# A wearable platform for closed-loop stimulation and recording of single-neuron and local field potential activity in freely-moving humans

**DOI:** 10.1101/2022.02.05.479253

**Authors:** Uros Topalovic, Sam Barclay, Chenkai Ling, Ahmed Alzuhair, Wenhao Yu, Vahagn Hokhikyan, Hariprasad Chandrakumar, Dejan Rozgic, Wenlong Jiang, Sina Basir-Kazeruni, Sabrina L. Maoz, Cory S. Inman, Jay Gill, Ausaf Bari, Aria Fallah, Dawn Eliashiv, Nader Pouratian, Itzhak Fried, Nanthia Suthana, Dejan Markovic

## Abstract

Advances in technologies that can record and stimulate deep-brain activity in humans have led to impactful discoveries within the field of neuroscience and contributed to the development of novel therapies for neurological and psychiatric disorders. Further progress, however, has been hindered by device limitations in that recording of single-neuron activity during freely-moving behaviors in humans has not been possible. Additionally, implantable neurostimulation devices, currently approved for human use, have limited stimulation programmability and lack full-duplex bi-directional capability. Here, we developed a wearable bi-directional closed-loop neuromodulation system (Neuro-stack) and used it to record single-neuron and local field potential activity during stationary and ambulatory behavior in humans. Together with a highly flexible and customizable stimulation capability, the Neuro-stack provides an opportunity to investigate the neurophysiological basis of disease, develop improved responsive neuromodulation therapies, explore brain function during naturalistic behaviors in humans, and consequently, bridge decades of neuroscientific findings across species.

## Introduction

Understanding brain function and its relation to cognition and behavior requires the integration of multiple levels of inquiry, ranging from the examination of single cells all the way up to the probing of human experience under naturalistic conditions. One major barrier that separates these approaches is the inability to record from single neurons during naturalistic behaviors in humans, which frequently involve full-body locomotion as well as twitches, gestures, and actions of the face and hands. This is problematic because behaviors that are studied in animal neurobiology are done almost exclusively in freely-moving animals (e.g., rodents) [1–2]. Thus, major gaps remain between understanding findings from neuroscience studies in animals to those in humans.

In parallel with progress in neuroscience, the medical field has seen a significant increase in the use and development of therapies delivered through implanted neural devices to treat and evaluate abnormal brain activity in patients with neurologic and psychiatric disorders [3–8]. However, current implantable devices do not allow for the recording of single-neuron activity, nor do they allow for extensive customization of stimulation parameters (e.g., pulse shape, precise timing with respect to ongoing neural activity), capabilities which would significantly expand the types of research questions that can be investigated. Furthermore, there is a critical need for robust data analytic capabilities on these devices (e.g., using deep learning and artificial intelligence) to deal with the large and complex neural data in real-time. Finally, an additional impediment in developing new responsive neurostimulation treatments is the lack of a customizable bi-directional interface that can record simultaneously with stimulation (full-duplex) and thus “talk” with the brain at the speed of behavior and cognition.

Since neural mechanisms underlying specific behaviors or brain disorders can span across a large population of cells, often from widespread brain regions [9, 10], there is a need for implantable neural devices to record from an increased number of channels across the brain. Further, there is a need for a sufficient temporal scale (<1 ms) to capture both single-neuron and local field potential (LFP) activity. Importantly, such technology should have a minimal impact on a person’s ability to move freely. Current neuroimaging techniques used in humans (e.g., functional magnetic resonance imaging [fMRI], scalp electroencephalography [EEG], magnetoencephalography [MEG]) have insufficient combined spatial and temporal resolution to record single-neuron activity. Intracranial electrophysiological studies, using micro-wire electrodes in epilepsy patients, can record LFPs and single-unit activity, however research participants must be tethered to large equipment and remain immobile. The high spatiotemporal resolution of LFPs (1–10 mm, ≥1 ms) and single-unit (10–50 μm, <1 ms) recordings comes at the cost of brain coverage, which is mitigated, whenever possible, with a larger number of recording channels through clinically-guided implantation of 10-15 depth electrodes (i.e., in stereo-EEG [SEEG]). In this realm, there are two possibilities for neuroscience studies to leverage clinical opportunities where individuals have electrodes implanted in their brains. The first is to use in-clinic research equipment (e.g., Blackrock Microsystems [11], Neuralynx [12], Nihon Kohden [13], Ripple Neuro [14]) with immobile participants undergoing clinically indicated SEEG who participate in voluntary research studies while hospitalized. Stimulation research studies are similarly done bedside, primarily using open-loop stimulation [15–21], although recent studies have begun to explore the use of closed-loop stimulation [22–26]. Critically, the equipment used in these research studies is expensive (up to ~$200K), bulky, and does not allow for extensive on-device customization of stimulation or complex real-time analyses for closed-loop stimulation. The second option is to use FDA-approved commercially available neural devices already implanted in several thousand individuals to treat epilepsy and movement disorders (e.g., Neuropace RNS System [27] and Medtronic Percept [28]). These chronically implanted devices offer research participants mobility at the expense of using large macro-recording electrodes that cannot record single-unit activity, fewer channels (usually 4 bipolar), and lower sampling rates (250 Hz). Other investigational devices such as the Medtronic Summit RC+S [29–31], allow for recording 16-channel intracranial EEG (iEEG) activity at up to 1 kHz sampling rates (no single-units). However, they are not FDA-approved for clinical treatment and thus exist in only a handful of patients with an FDA investigational device exemption (IDE) approval, limiting their widespread use by the scientific community. Existing closed-loop implantable technologies also lack full-duplex ability, which allows for simultaneous stimulation and recording of neural tissue inclusive of unit and LFP activity. While research studies using these systems have given rise to several impactful neuroscientific discoveries [27, 32], the possibility of novel devices to one-day record from single neurons, deliver customizable closed-loop stimulation, and carry out complex data analytics in real-time would provide unparalleled opportunities for first-in-human scientific discovery and the development of more effective medical therapies for patients’ neuropsychiatric disorders.

Here, we present a potential technological pathway towards future more advanced implantable technologies with the development of a miniaturized bi-directional neuromodulation external device (Neuro-stack) that can record up to 256-channel (128 monopolar/bipolar macro-recordings) iEEG and 32-channel single-unit/LFP activity from micro-wires during ambulatory behaviors in humans who have macro- and micro-wire depth electrodes implanted for clinical reasons. The Neuro-stack can deliver customizable closed-loop multi-channel (up to 32 simultaneous) stimulation where parameters such as pulse shape, frequency, amplitude, pulse width, inter-pulse width, polarity, channel selection and timing (e.g., for phase-locked stimulation) are configurable. A major advantage of the Neuro-stack is its full-duplex capability that allows for the recording of neural activity in the presence of concurrent stimulation.

We include data acquired using the Neuro-stack showing single-unit, LFP, and iEEG activity recorded in twelve participants who had depth electrodes implanted for epilepsy evaluation. In one of these participants, we used the Neuro-stack to perform binary prediction of memory performance in real-time (69% F1-score) using neural activity recorded from medial temporal lobe (MTL) regions. We also demonstrated the Neuro-stack’s ability to record single-neuron activity during walking behavior and deliver customized stimulation. These capabilities can be useful for future studies investigating the neural mechanisms underlying naturalistic behaviors in humans and developing novel neuromodulation therapies for patients with brain disorders that will be effective in real-world settings.

## Results

The Neuro-stack (Fig. 1a-b) provides a bi-directional neuromodulation platform for wide-band sensing and stimulation of deep-brain areas for basic and clinical neuroscience studies. Compared to much larger existing devices (Fig. S1) that are used bedside and carried on a cart, the Neuro-stack’s small hand-held size enables concurrent stimulation and recording of real-time electrophysiology (single-unit and LFP activity) during freely-moving behavior (Fig. 2) by connecting to commonly used implanted macro- and micro-electrodes (Fig. 1c-d). Apart from its small form-factor and unique on-body wearability, the Neuro-stack can support:

1. *Recording* of up to 256 channels for a total of 128 monopolar or bipolar recordings with a sampling rate of up to 6,250 Hz. Further, wide-band sensing from up to 32 monopolar or bipolar recordings at up to 38.6 kHz allows for the recording of single-unit and LFP activity simultaneously.
2. *Flexible and programmable stimulation* (Fig. 3) allowing for delivery of bipolar/monopolar stimulation to any 32 out of 256 contacts simultaneously. Stimulation engines are current-controlled and allow the user to program current amplitude, frequency, timing, pulse shape, and other parameters (Fig. 3, Table S2).
3. *Closed-loop neuromodulation*. The Neuro-stack has built-in (hardware) oscillation power detection and thus the ability to trigger stimulation at a predefined phase of an oscillation (phase-locked stimulation [PLS] delivered at a particular phase of ongoing theta activity). Further, sensing of neural activity is concurrent with stimulation for true (full-duplex) closed-loop capabilities. Resources for designing custom closed-loop algorithms are available at both the embedded hardware and external software levels.
4. *Software support* that comes in two formats. First, a turnkey graphical user interface (GUI) running on a Windows-based tablet or laptop is available for research purposes (Fig. 1a). Second, a full-access application programming interface (API) library written in C++ allows the user to build custom research open- and closed-loop stimulation capabilities for research studies (Fig. S3).
5. *Tensor multiplication accelerator* (Edge TPU, Fig. 2a, Fig. S3 middle) that is integrated with the Neuro-stack device, enabling an extended range of applications such as real-time inference for neural decoding (Fig. 4) or closed-loop stimulation.
6. *Wired or wireless mode*. The Neuro-stack platform can be externally controlled and powered via a USB cable or remotely controlled through a secure local network using a battery-powered configuration (Fig. 2a, Fig. S3). This flexibility allows researchers to perform wide-band recording and stimulation during either stationary or ambulatory (freely-moving) behavioral tasks.

**Figure 1.**
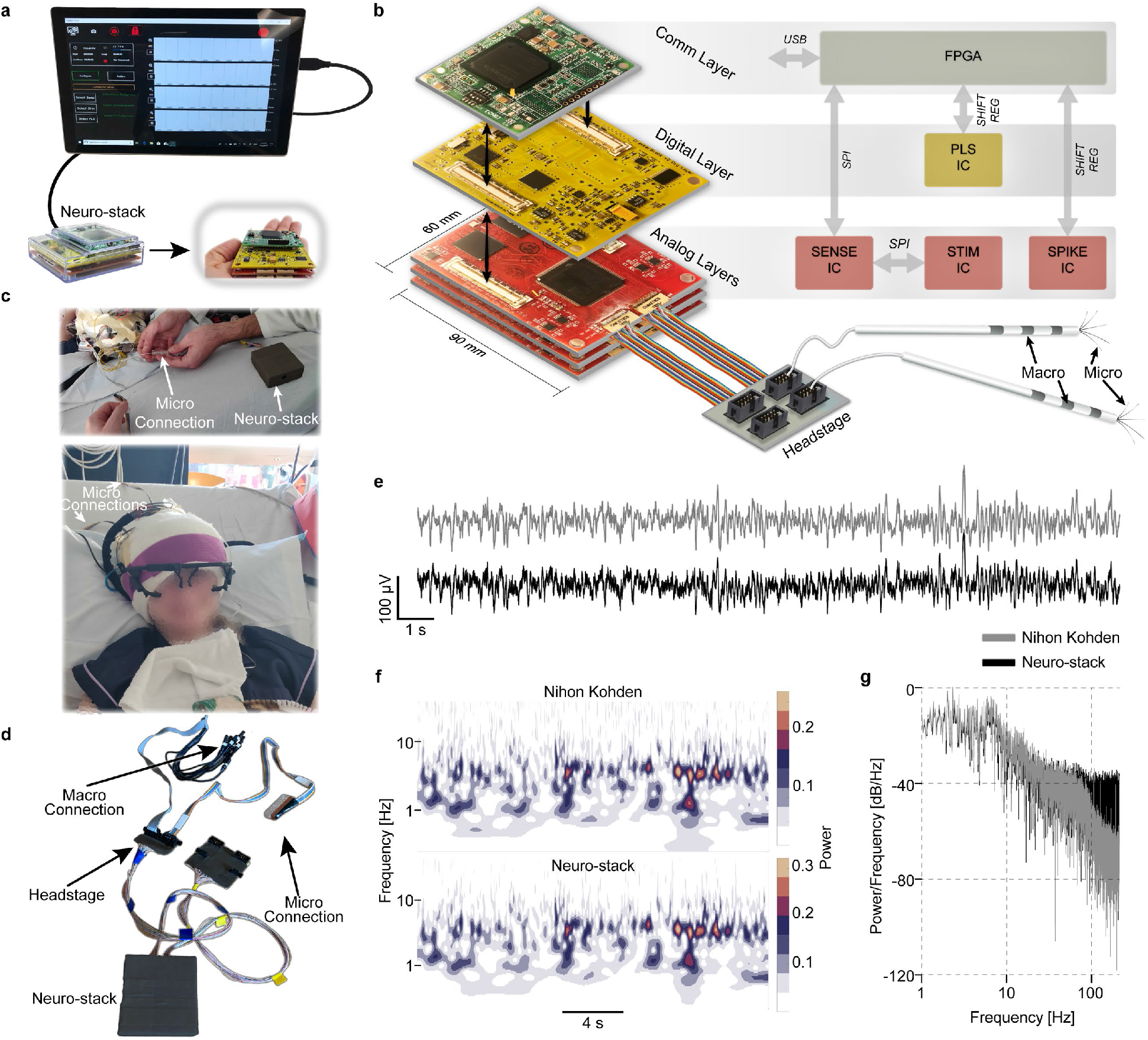
Neuro-stack platform. **a,** Neuro-stack and GUI-based tablet for single-neuron and local field potential (LFP) recordings, and closed-loop programmable phase-locked (PLS) stimulation. The tablet allows for selection of recording and stimulation channel(s), sampling rate, monopolar/bipolar recordings, and other parameters. Shown are the packaged (left) and unpackaged (right) versions. **b,** The Neuro-stack consists of three stacked layers: 1) Communication (Comm), 2) Digital, and 3) Analog. Presented are the printed circuit boards (PCBs, size = 90×60 mm^2^) and 5×2 pin (8 channels, 1 reference, and 1 ground, 10 total pins) Omnetics headstage connectors to which micro-electrodes can be connected (only top Analog layer connected). Note that each Analog layer receives up to two Omnetics connectors to connect with up to 4 electrodes through one headstage. A high-level block diagram of each layer is shown (right). The Comm Layer contains a FPGA (field-programmable gate array) that mediates command and data transmission (via USB) between external software and integrated circuit (IC) chips. The Digital Layer contains the PLS IC. The Analog Layer contains chips for sensing (Sense IC) and stimulation (Stim IC). Three Analog layers are shown to allow recording of 192 channels (64 x 3 layers). Serial peripheral interface (SPI) is used for FPGA communication with the Sense and Stim ICs, and shift register for FPGA communication with the PLS and Spike ICs. **c,** The Neuro-stack connected to micro-electrodes in a participant wearing an eye-tracking system. **d,** Shown are 10-pin touch proof jumpers for macro-electrode and 10-pin connectors (e.g., Adtech) for micro-electrode recordings. **e,** Example data recorded simultaneously using a clinical monitoring system (Nihon Kohden, gray) and Neuro-stack (black) showing similarity of signals. **f,** Example power spectrograms from data (**e**) showing concordant activity patterns. Frequency (0.1–32 Hz) is shown using a logarithmic scale. **g,** Example normalized power spectral density (PSD) plots from data shown in **e**.

**Figure 2.**
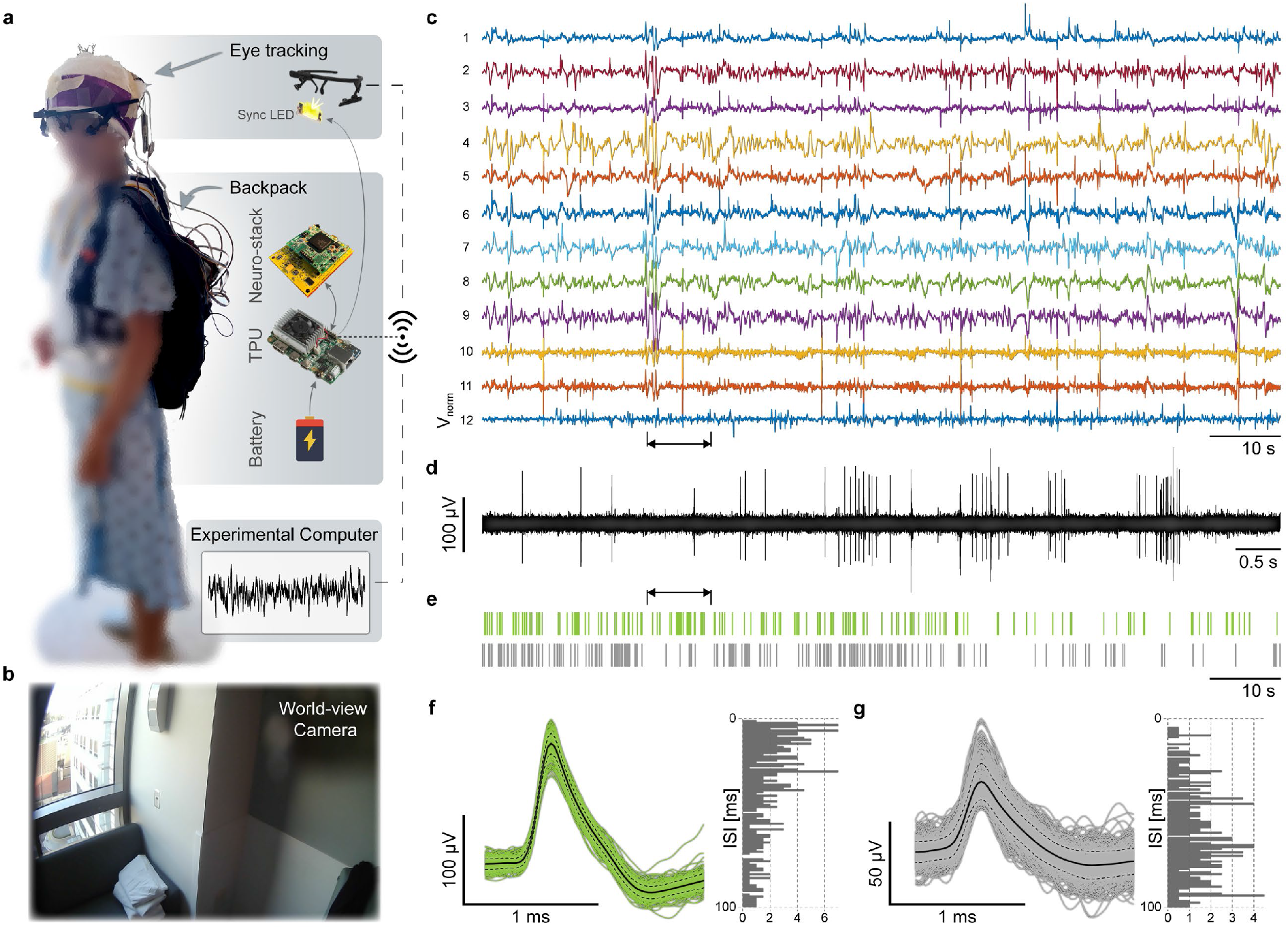
Neuro-stack as a wearable platform for recording neural activity during ambulatory behavior in humans. **a,** An example research participant wearing the backpack carrying the Neuro-stack system, a single board computer with a tensor processing unit (TPU), and a battery, to allow for recording of single-neuron and LFP activity during ambulatory behavior. The participant was also wearing an eye-tracking device that keeps track of head direction, pupil size changes, eye movements. Data captured from the eye-tracker was synchronized with the neural data using a programmable light emitting device (LED) that is visible on the eye-tracker world-view camera. Wireless communication between the Neuro-stack, eye-tracker, and other external monitoring devices is enabled through a Wi-Fi access point on the TPU device. **b,** Neural activity was recorded during an ambulatory task where participants walked repeatedly (10 times) between two opposite corners of a 5 x 5 ft^2^ room (from X to Y, Fig.S2b). Example video frame from the eye-tracking world-view camera as an example participant approached point Y in the room (bottom). **c,** Neural activity (voltage-normalized separately for each channel) from 12 micro-electrode channels (1-6: hippocampus, 6-12: anterior cingulate) during the ambulatory walking task from an example participant. **d,** 10 s of filtered data from channel 12 (arrows point to corresponding sections on **c** and **e**). **e,** A raster plot of two single-units isolated from channel 12. **f**, The first single-unit isolated from channel 12 and its corresponding inter-spike interval (ISI) histogram (right) **g,** The second single-unit isolated from channel 12 and its corresponding inter-spike interval (ISI) histogram (right).

**Figure 3.**
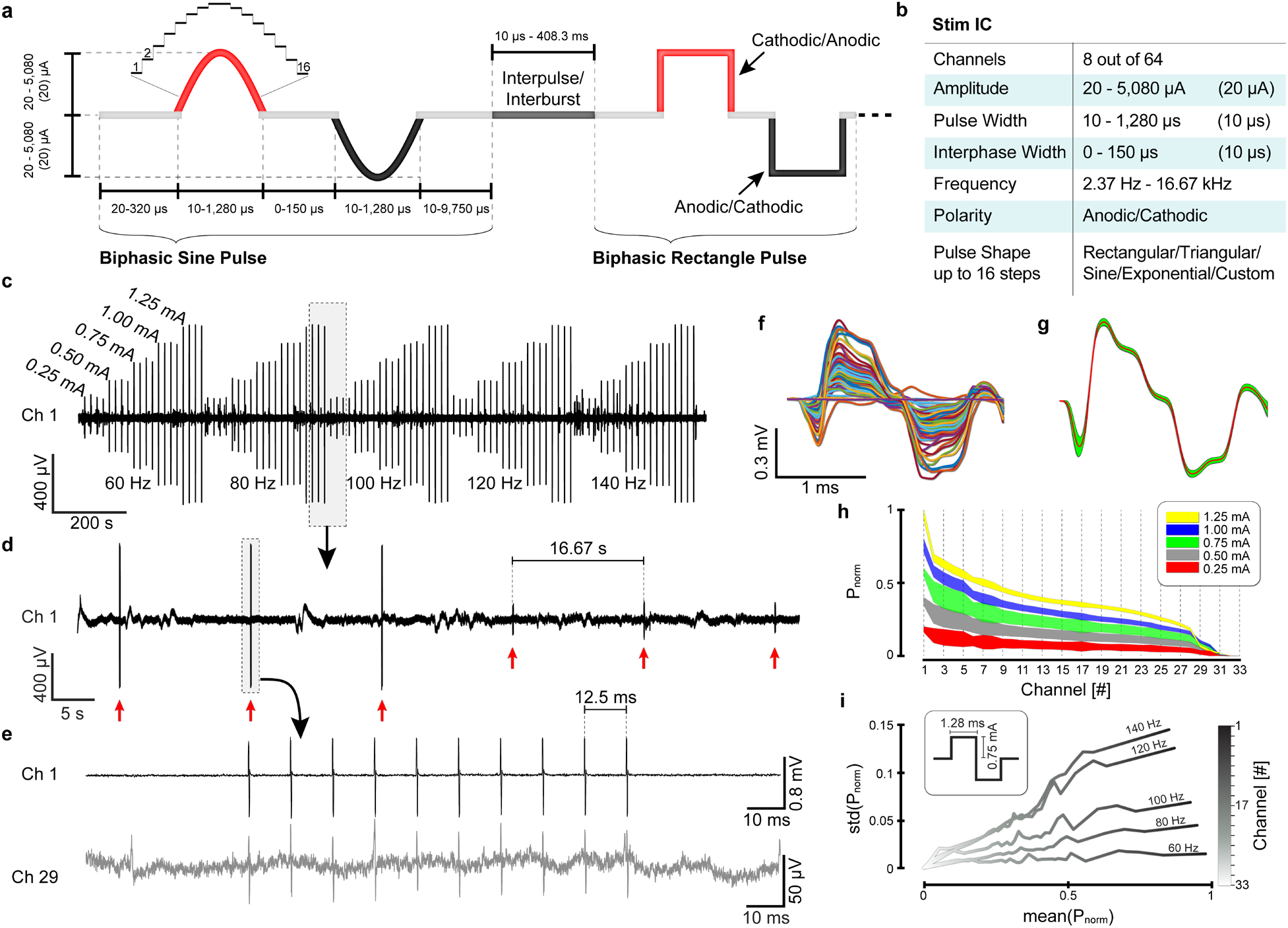
Neuro-stack as a programmable closed-loop neuromodulation system. **a,** Stimulation parameters can be customized including frequency, amplitude (0-5080 mA in steps of 20 mA), polarity (anodic/cathodic), timing (of pulse width, inter-phase width and inter-pulse/burst interval) and pulse shape (e.g., sinusoidal or rectangular pulses shown). **b,** Key features and capabilities on the stimulation integrated circuit (Stim IC) including the number of channels (i.e., 8 out of 64 per analog layer) that can be selected for stimulation, amplitude, configurable pulse shapes where amplitude in each of up to 16 steps (**a**) can be programmed for custom waveform design, frequency, polarity, pulse width (**a**, 10-1280 μs, steps: 10 μs), and inter-phase width (**a**, 0-150 μs, steps: 10 μs). **c,** Example macro-electrode channel recorded during the delivery of macro-stimulation, which was delivered with varying combinations of amplitudes × frequencies [(0.25, 0.5, 0.75, 1.00, 1.25) mA × (60, 80, 100, 120, 140) Hz]. Each stimulation burst contained 10 biphasic (pulse width = 1.28 ms) after which a delay of 16.67 seconds occurred before the next burst cycle. **d,** Zoomed-in view of **c** (outlined box) where six stimulation bursts (red arrows) are shown with different parameters (burst 1-3: 1.25 mA, 80 Hz; burst 4-6: 0.25 mA, 100 Hz). **e,** Zoomed-in view of a single burst (outlined box) from the same channel in **d** and another example channel (29). **f,** Time-aligned bipolar pulses from a stimulation burst (10 pulses, 1.25 mA) from all channels (n=33). **g,** Mean and standard deviation (std) values of all time-aligned bipolar stimulation pulses from example recording channel (1). **h,** Normalized power (mean and std) of the propagated stimulation pulses across channels (n=33) recorded with respect to varying stimulation current (0.25–1.25 mA). **i,** Std of normalized power (std(power/max[power])) as a function of mean normalized power (mean(power/max[power])) differentiates pulse propagation across channels with respect to varying stimulation burst frequencies (60-140 Hz, steps: 20 Hz) with a fixed pulse width (1.28 ms) and current amplitude (0.75 mA). Electrode channels are marked in shades of gray (n=33).

**Figure 4.**
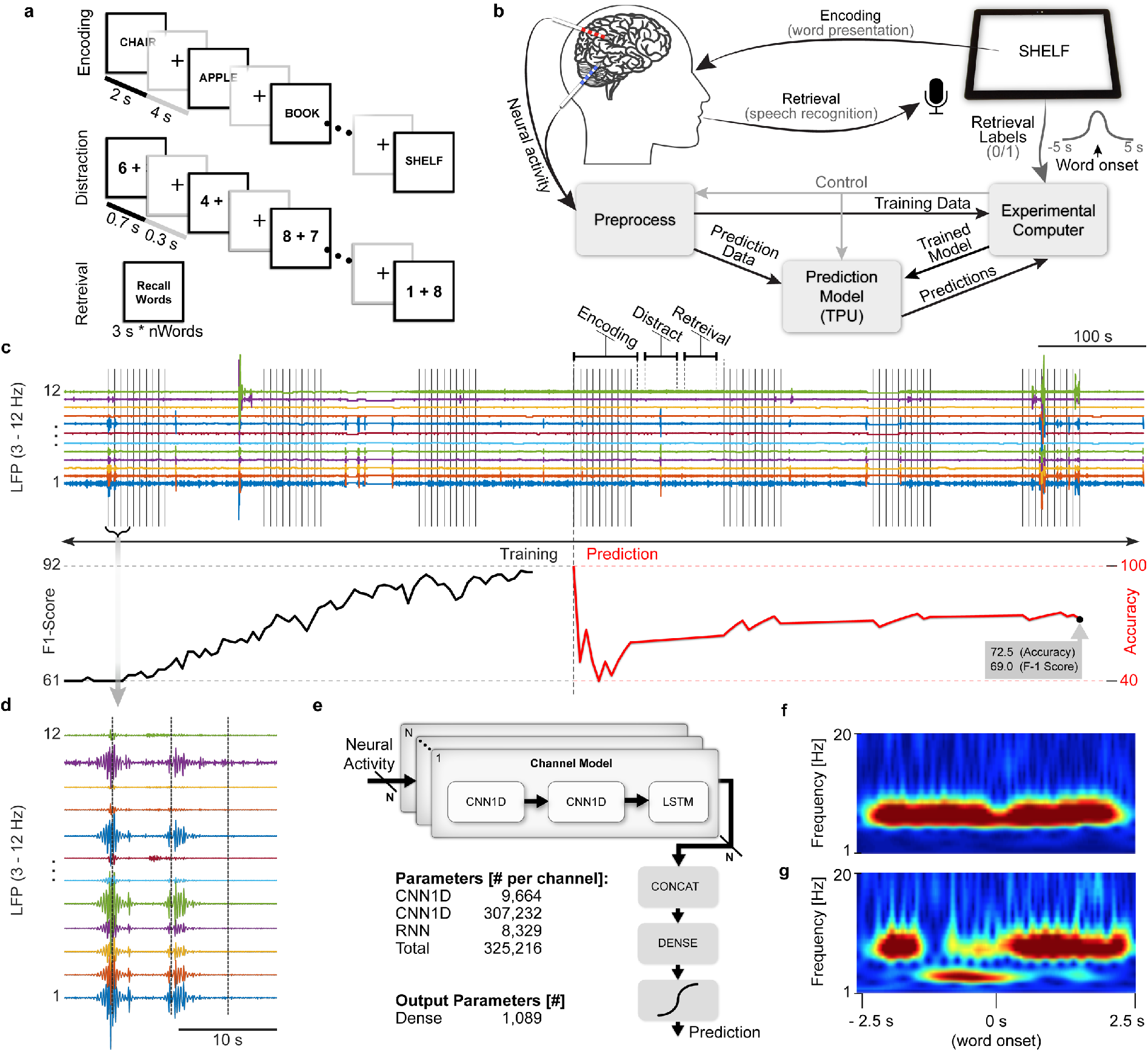
Decoding memory performance with the Neuro-stack system. **a,** Neural activity was recorded during completion of a verbal memory task, which included three phases: 1) Learning (encoding) during which a list of words presented (2 s each, 0.8 s inter-stimulus interval [ISI]), 2) Distraction, during which numbers were presented serially (0.7 s each, 0.3 s ISI) and participants were instructed to respond odd/even, and 3) Recall (retrieval) where previously presented words were recalled. **b,** Neuro-stack recording setup and processing pipelines used during the memory task. A tablet was used to present words during Encoding and record to identify in real-time the spoken words recalled during Retrieval (using speech recognition). Minimally processed data was then fed into an external computer with synchronized retrieval results. The neural network model (Model, **e**) was trained in real-time to predict retrieval performance based on neural activity during encoding. The model was then ported to the TPU to perform real-time predictions. **c,** Filtered theta (3–12 Hz) activity from the left hippocampus (LHC) is shown since it was the most critical feature used by the trained neural network model to predict memory (top). Vertical lines mark the onset of each word (10) during 7 repetitions (blocks) shown of the memory task. Decoding performance (accuracy) is shown (bottom) during the first three blocks, which were used to train the neural network (Training) and the associated aligned F1-Score. The last four blocks were used to predict memory performance (Predict) and the associated aligned F1-Score. **d**, Zoomed-in-view of example theta activity shown in **c**. **e,** The neural network model (2 × CNN1D + LSTM + Dense network) parameters. **f,** Time-frequency representation of the first most significant feature (from the trained CNN layer activation filter), which highlights theta power during encoding. **g**, Time-frequency representation of the second most significant feature (trained CNN layer activation filter), which highlights temporal patterns in theta activity with increases particularly after the onset of word presentation.

The central hardware component of the Neuro-stack platform (Fig. 1a-b) consists of three printed circuit board (PCB) layers: 1) analog, 2) digital, and 3) communication. Each layer is embedded with one or several dedicated integrated circuit (IC) chips. The analog layer (Fig. 1b, bottom) contains mixed-signal sensing IC (Sense IC and Spike IC) and stimulation IC (Stim IC) chips, which were previously developed as part of the DARPA SUBNETS program [33–36]. A single Sense IC (one per analog layer) accepts neural activity from up to 64 electrode contacts fed into voltage-controlled oscillators (VCO), which serve as analog-digital converters (ADC). Each VCO ADC supports 6,250/N Hz sampling frequencies, where N = 1,2,4,8,…, 128 and a 100 mV_pp_ linear input dynamic range with 12/21 (macro/micro) bits of resolution, ensuring that the underlying neural signal is captured in the presence of large artifacts (e.g., from stimulation). The Sense IC contains digital nonlinearity correction to account for nonlinear amplification across the input range. Moreover, it also contains a digital logic for adaptive stimulation artifact rejection that subtracts a template stimulation artifact extracted from adjacent channels [36]. The total power consumption per channel is 8.2 μW. A single Spike IC (one per analog layer) accepts neural activity from up to 8 micro-wire contacts and supports sampling rates of up to 38.6 kHz [37]. A single Stim IC contains eight engines that can, with the appropriate configuration, drive current through any individual or combination of the connected 64 electrode contacts. Stimulation output current is highly configurable (Fig. 3a, b), including selection of amplitude, frequency, and multiple or custom waveform shapes. This flexible programmability allows for stimulation using previously used burst protocols [38–40] as well as exploration of novel stimulation patterns for investigative research and therapy development. These capabilities also enable increased degrees of freedom (timing, amplitude parameters; Fig. 3) compared to currently available intracranial neurostimulation systems (Table S2).

The Neuro-stack’s digital layer (Fig. 1a-b, middle) routes signals between the analog and communication layers and contains a custom IC chip (PLS IC) for closed-loop stimulation based on the detected oscillatory (e.g., theta) phase in the recorded neural signal coming from the analog layer to enable PLS [41–42]. A field-programmable gate array (FPGA, Xilinx Spartan 6 board) serves as a communication layer (Fig. 1a-b, top, Fig. S3) between an external devices and custom ICs (Fig. 1b).

The Neuro-stack uses the serial peripheral interface (SPI) at 12 MHz (Sense IC and Stim IC) and serial shift register (PLS IC and Spike IC) for internal communication between layers and IC chips and a USB interface for external communication and power supply. The device is assembled by physically stacking the described layers (Fig. 1a-b). Furthermore, one Neuro-stack device supports up to four analog layers at the same time, for up to 256 micro-wire (LFP) electrode contacts (64 per layer) and up to 32 micro-wire (single-unit) electrode contacts (8 per layer).

A ready-to-use GUI is available (connected to the Neuro-stack via USB) and allows for real-time multi-channel monitoring and control of neural recording and stimulation (Fig. 1a). A platform-agnostic API library written in C++ that allows for custom applications and experiments is also provided. To allow ambulatory experiments, the Neuro-stack can be wirelessly controlled using the Coral Development Board (CDB; Fig. S3, TPU in Fig. 2a), an ARM-based single-board computer, running a Mendel Linux distribution. Similar microprocessors with wireless capabilities such as a Raspberry Pi can also be used for this purpose. Our Neuro-stack setup included an ARM-compiled Neuro-stack API, which supports wireless applications through a secure local Wi-Fi (2.4 or 5 GHz) network created using the included API library. Only an experimental device that uses a secure (X.509 certification) connection to a local server can control the Neuro-stack. The CDB contains an onboard TPU (Fig. 2a), which can make real-time inferences for neural decoding or closed-loop applications (e.g., Stationary Verbal Memory Task section, Fig. 4).

### In-vitro Sensing and Stimulation

The Neuro-stack IC chips (i.e., Stim, Sense, Spike, PLS) were validated in-vitro separately [33–34, 36–37, 41] and some (Sense and Stim) as part of an implantable system [34]. Before moving to human in-vivo studies, in-vitro validation of all chips in the Neuro-stack was also completed. The setup for validating sensing capability included the feeding of pre-recorded analog neural data via an NI PXI System (digital to analog converter) through a phosphate-buffered saline (PBS) solution, use of an oscilloscope to observe true signals at front-end inputs, and a computer to control and power the Neuro-stack (Fig. S4, Online Methods). The captured signals were of satisfactory quality (Sense and Spike IC, Fig. S5a,b).

PLS was also tested using the same in-vitro setup (Fig. S4). For 300 s of LFP data, the results showed 400 detections within the theta band (3–8 Hz) and triggered stimulations with a circular variance of 0.3 [42].

Measurements of stimulation and synchronization delivery delays were also characterized for ensuring accurate closed-loop implementation as well as alignment between behavioral stimuli, neural data, and other devices that run in parallel. First, the round-trip delay, important for closed-loop stimulation, was measured from sensed input to stimulation output by feeding a train of 50 pulses into the sensing front-end. The pulse rising edge detection triggered stimulation on the CDB software side (connected to the Neuro-stack via USB; Fig. S5c). Input/output observations by the oscilloscope showed a 1.57 ± 0.19 ms round-trip delay (Fig. S5d). This result was consistent with the PLS-based round-trip delay of 1.7 ± 0.3 ms measured from the sensed input to stimulation output [42]. Second, synchronization with external devices was done by timestamping neural samples using the CDB; accuracy depended on the system latency through hardware and software. We applied the same approach as the round-trip delay with the addition of sending a test pulse on a general-purpose pin once the sample reached the timestamping step (Fig. S5c), which resulted in a delay, measured from sensed input to CDB output, of 0.56 ± 0.07 ms (Fig. S5e). For more details see Online Methods – Neuro-stack in-vitro testing section.

### In-vivo Sensing and Stimulation

Twelve participants with indwelling macro- and micro-wire electrodes implanted for pharmacoresistant epilepsy volunteered for the study by providing informed consent according to a University of California, Los Angeles (UCLA) medical institutional review board (IRB) approved protocol. Each Behnke-Fried macro-micro depth electrode (Ad-Tech Medical, Racine, WI) contained 7-8 macro-contacts and 9 (8 recording, 1 reference) 40-μm diameter platinum-iridium microwires [43] inserted through the macro-electrode’s hollow lumen. Neural activity was recorded from macro- and micro-wire contacts using the Neuro-stack during wakeful rest in all participants (Online Methods - Participants) and from various brain regions (Table S1; Online Methods – Electrode Localization). The Neuro-stack setup was done bedside (Fig. 1c-d) or on-body during ambulatory movement (Fig. 2a), where the system was connected to implanted electrodes using a custom-built connector (i.e., touch-proof, Cabrio, and Tech-Attach connectors for commercial Behnke-Fried macro- and micro-electrodes, respectively). The main objective of the in-vivo validation studies was to test recording of single-unit and LFP activity and macro-stimulation during rest and behavioral tasks (see Ambulatory Walking Task and Stationary Verbal Memory Task sections). The PLS closed-loop functionality has been tested in-vitro [42] with expected in-vivo validation to be a part of future behavioral studies.

iEEG data was also recorded simultaneously with the Neuro-stack using commercially available electrophysiological recording systems (i.e., Nihon Kohden) for comparison purposes. Example raw iEEG activity traces from one participant is shown using simultaneous Nihon Kohden and Neuro-stack recordings (Fig. 1e), together with time-frequency power spectrum data (frequency band: 1 – 32 Hz; Fig. 1f), and PSD plots (frequency band 1 – 250 Hz; Fig. 1g).

Stimulation was performed in three participants to test stimulation artifact propagation across channels and assess associated statistics with varying parameters. In the first two participants, bipolar macro-stimulation was applied to the left hippocampus (amplitude: 0.5 mA; Pulses/burst: 11; waveform shape: rectangular; pulse width: 1ms; frequency: 100 Hz). After successful delivery was observed in surrounding channels, a series of bipolar macro-stimulation bursts with varying parameters was delivered in a third participant. The parameter test space included [amplitude, frequency] combinations of [0.25, 0.50, 0.75, 1.00, 1.25] mA × [60, 80, 100, 120, 140] Hz where every combination was repeated four times for a total of 100 macro-stimulation bursts (Fig. 3c) with the following parameters (pulse width: 1.28 ms, interphase width: 150 μs, rectangular pulse shape, interburst delay: 16.67 s). Stimulation delivery (Fig. 3c–entire session; Fig. 3d–multi burst; Fig 3e–single burst level) was observed on 40 nearby recording channels, obtained using the Sense IC (sampling rate: 6250 Hz). Overlayed pulses from one of the bursts with the same parameters (1.25 mA, 60 Hz) showed successful delivery across all channels (Fig. 3f – upsampled to 25 kHz and interpolated). Further, all pulses from the same burst showed consistent artifacts in the channel adjacent to the stimulation site (Fig. 3g – mean ± std). Higher stimulation amplitudes resulted in lower variability (std) in delivered power (Fig. 3h) while higher burst frequency resulted in higher variability across channels (Fig. 3i). Note, that stimulation artifacts were not caused or affected by channel saturation (Fig. 3f) with absolute voltage levels much lower than the 50 mV cut-off. Results (Fig. 3h,i) suggest that deviation of underlying neural activity is not the only cause of artifact waveform uncertainty. Future studies that model and predict artifact propagation could use stimulation mapping prior to studies to characterize effects and adjust expected values and deviations accordingly.

### Ambulatory Walking Task

We used the wireless Neuro-stack setup (Fig. 2a) in six participants while they walked in their hospital rooms to record single-neuron activity from various brain regions (Table S1) synchronized with world-view and eye-tracking cameras (Fig. 2b, Fig. S2c-d, Online Methods).

The first four participants walked freely around the room, during which motion artifacts in recordings were examined. The use of nearby electrodes (same bundle) as a reference resulted in reduced common noise artifacts using the front-end amplifiers (Fig. S2a). The last two participants were walked from one point of the room to another ten times (Fig. S2b). Raw (line noise removed) 12-channel neural activity recorded from one participant during walking is shown in Figure 2c. Although motion artifacts were reduced, slow voltage transients during movement were still present (Fig 2c). Nonetheless, single-unit spikes were preserved (Fig. S2a-right) and detected using a bandpass filter [300 – 3000 Hz] (Fig. 2d, Fig. S2a-right). After spike sorting [44] of the data, single-unit clusters were successfully isolated (Fig. 2e-g).

### Stationary Verbal Memory Task

Neuro-stack’s ability to record neural data in real-time and decode behavioral performance was tested bedside in a participant with indwelling micro-wire electrodes while they completed a verbal memory task (Fig. 4a). During the task, the participant was instructed to learn (encode) a list of ten words that were presented on an iPad screen and then verbally recall as many words as possible after a brief delay (30 s). During the delay, a non-mnemonic (distraction) task was completed that involved identifying whether the sum of the two random numbers (1-9) was either odd or even. Encoding, distraction, and recall blocks were repeated nine times during the experimental paradigm while the Neuro-stack recorded LFP activity from sixteen micro-wire channels, which was used to decode memory performance in real-time using artificial neural networks.

The TPU device (Fig. S3) was integrated with the Neuro-stack and used to embed a neural network model that was large enough to generalize across participants but small enough to be successful with using solely on-system computation. Artificial neural networks were pre-trained on multi-channel raw (downsampled) LFP data previously acquired using a Blackrock Neuroport recording system. Offline pre-training performance successfully differentiated remembered from forgotten words during recall with a test F1 score (*F*_1_ = 2 × (*P* × *R*) / (*P* + *R*), *P* – precision, *R* – recall) of 88.6 ± 5.5% and a test accuracy of 91.7 ± 3.3%. The model was built and trained in a Keras (TensorFlow backend) framework after detailed comparison with commonly used machine learning methods (Support Vector Machine [SVM], Principal Component Analysis [PCA] plus SVM, various neural network architectures; Table S3). The decoder consisted of an input 2 × CNN1D + LSTM layers that extracted multi-channel LFP features and an output Dense (fully connected network) + Classifier layers (Fig. 4e). For further details see Online Methods.

During the memory task, the offline model’s output layers were retrained in real-time on an external computer. The trained model was then translated to TensorFlow Lite and ported to the Edge TPU, to predict memory during the last four task blocks (Fig. 4b). The training phase and improving accuracy/loss metrics for an example participant are presented in Fig. 4c. The online test (prediction) phase resulted in an F1 Score of 69% (Fig. 4c-bottom). Average total power at theta frequency bands (Fig. 4d) indicated a significant difference between correct and missed trials. A time-frequency heatmap of the second CNN1D layer activation filters [45] confirmed that theta multimodal activity timed to the population activations in the left and right hippocampus was used by the model to identify correctly recalled words (Fig. 4f, g).

## Discussion

We present the Neuro-stack, a novel miniaturized recording and stimulation system that can interface with implanted electrodes in humans during stationary (bedside) or ambulatory behaviors. The Neuro-stack can record up to 256 channels of LFP/iEEG activity and 32 channels of single-/multi-unit activity. Macro-stimulation can also be delivered through any of the channels (up to 32 channels simultaneously) during recording, allowing for bi-directional full-duplex capability. This is a significant advantage over existing systems in that it allows for characterization of ongoing neural consequences of stimulation as well as precisely timed closed-loop stimulation.

A second major advantage of the Neuro-stack over existing systems is its smaller hand-held size that enables it to be carried on-body and be wirelessly controlled. These features allowed us to record single-neuron waveforms (spikes) during walking, which to our knowledge are the first recordings of their kind in humans. Future studies using the Neuro-stack could determine the neural mechanisms underlying human freely-moving behaviors (e.g., spatial navigation) to identify, for example, spatially selective neurons and their modulation by cognition (e.g., hippocampal place or entorhinal grid cells [46]) that have been previously discovered in freely-moving animals. Doing so would bridge decades of findings between animals and humans and potentially lead the way towards scientifically informed therapies for hippocampal-entorhinal-related dysfunctions (such as Alzheimer’s disease). While we did not identify any spatially selective single-units in the current study, possibly due to the restricted spatial environment in which walking took place, further analysis from our ambulatory task and other future studies using the Neuro-stack over longer distances (e.g., hallways) may be able to identify these neurons in humans.

A third advantage of the Neuro-stack is its API that allows fast and flexible prototyping of the experiments with range of backend functions that accurately align behavioral and neural events (i.e., spikes). We demonstrated how the Neuro-stack’s API integrated with a TPU can, in real-time, decode verbal memory performance in a single participant with accuracy levels that are comparable to previous reports [15]. Specifically, we used neural network models applied to hippocampal recordings to predict whether a previously learned item would be remembered, with offline results exceeding those previously reported [15], when equivalent metrics (F1-scores at the optimal thresholds) are compared. Future studies with larger sample sizes will confirm whether reported decoding accuracy can be generalized across subjects. It should be noted that we tested the decoding algorithm in one participant using the model pretrained with recordings from a different device with different noise levels (Fig. 1g), hence it is reasonable to assume that performance could go up as more Neuro-stack data are incorporated into the pretrained model. Given the increasing benefit of using machine learning approaches [47–49] in neuroscience studies, the Neuro-stack could be useful for validating decoding models and testing novel closed-loop stimulation therapies (e.g., to improve memory in patients with severe memory impairments).

Future studies can also determine which stimulation parameters are most beneficial for restoring cognitive or behavioral functions given the Neuro-stack’s highly flexible programmability compared to existing human-approved stimulators. For example, continuous adjustments of custom pulse shapes, timing of complex burst patterns, and/or timing of stimulation relative to ongoing neural activity events could allow for the development of more effective stimulation therapies. Given the wireless and wearable nature of the Neuro-stack, studies could also determine whether closed-loop stimulation protocols effectively translate to more naturalistic behaviors during everyday experiences that occur during mobility.

While the Neuro-stack offers several advantages over currently available systems, there are limitations that warrant discussion. First, this Neuro-stack prototype can only support a maximum of 32 wide-band single-unit recording channels. While it can also simultaneously record up to 256 LFP recording channels (using four analog layers), other existing bedside systems can allocate more than 256 channels solely for unit recordings. The use of multiple Neuro-stack devices, however, would address this issue and increase single-unit channel count substantially. Second, although the Neuro-stack is small enough to be carried on-body and thus allow for full mobility, its connection with implanted electrodes is still wired, similar to other bedside systems. Thus, significant movements can result in motion artifacts. However, single-unit spike waveforms can still be detected and isolated during walking behavior as we show using techniques such as differential recordings between nearby contacts, as well as proper wire isolation and fixation. Lastly, the Neuro-stack currently can only be used in research studies with patients who have externalized electrodes implanted during clinical (e.g., epilepsy) monitoring. Since these patients need to be continuously tethered to bedside intracranial recording systems to assess for symptomatic episodes (e.g., seizures), this limits the amount of time a patient can be freely-moving. However, future studies can complete ambulatory studies after clinical data has been captured as was done in the current study, on the last day of the patient’s hospital stay prior to electrode de-plantation surgery, or during circumstances where continuous monitoring may not be necessary (e.g., depression or chronic pain studies [51–52]). Furthermore, proper precautions and safety measures should be implemented, such as waiting to complete studies until epilepsy patients are back on anti-epileptic medications to minimize risks associated with seizures during ambulatory tasks.

Although Neuro-stack is much smaller than other external systems, an even smaller version could be tested in future in-vivo studies since its IC chips are all implantable by design [29–33, 37] and require a combined area of just 113 mm^2^ (4 analog layers). An implantable version of the Neuro-stack [30] but with its added single-neuron and closed-loop stimulation capabilities thus presents an exciting avenue towards a completely wireless intracranial single-unit and LFP recording system that would not be susceptible to motion artifacts. This type of system would present a significant advancement over current FDA-approved chronic neurostimulation devices in that it would allow for single-neuron and multi-channel (current state-of-the-art is 4 channels; Neuropace RNS) recordings, bi-directional recording and stimulation (full-duplex) capability, and the ability to use advanced strategies for decoding (e.g., neural network models for inference) behavior or disease-related states. Altogether, these novel capabilities would provide cognitive and clinical neuroscience studies with a promising future pathway towards determining the deep-brain mechanisms of naturalistic behavior in humans and developing more effective closed-loop intracranial neuromodulation strategies for individuals with debilitating brain disorders.

## Contributions

Conceptualization: D.M., N.S., I.F., N.P., U.T., and S.B.; Methodology and Software: U.T., S.B., C.L., A.A., W.Y., V.H., H.C, D.R., W.J., and S.B-K; Investigation and data acquisition: U.T., S.B., C.L., C.S.I., J.G., and S.L.M.; Resources: A.F., A.B., D.E., I.F., N.S., and D.M.; Writing – Original Draft: U.T., N.S., and D.M.; Writing – Review and Editing: all authors; Visualization: U.T.; Funding: I.F., N.S. and D.M.

## Acknowledgments

This work was supported by DARPA SUBNETS (to D.M.), DARPA RAM (to IF and DM), the McKnight Foundation, NIH grants UO1 NS103802 and NS103780 (to N.S.), F30MH125534 (to S.L.M.), and NIH grant RO1 NS084017 (to IF). The authors thank Sonja Hiller, Edward Chang, Natalie Cherry, Guldamla Kalender, Andreina M. Hampton, Anthony J. Rangel, and Omar Morales for helpful discussions and assistance in the Neuro-stack design and testing. The authors also thank the participants for taking part in the in-vivo validation studies.

## Online Methods

### Neuro-stack Hardware Design

Neuro-stack was built from four implantable and previously reported application-specific integrated circuit (IC) chips. The Sense IC contains 32 low-noise, high dynamic range LFP sensing front-ends (FEs), which can be duplexed to 32 electrodes for single-ended recording with respect to the reference electrode or to 32 pairs of electrodes for differential recording, matching up to 64-electrode probe (or 8 × 10-electrode probes, where the 9^th^ is a reference and the 10^th^ a ground contact). After linearization in the nonlinearity correction (NLC) module, the recorded output can be optionally sent to 4 adaptive stimulation artifact rejection (ASAR) engines, which suppress stimulation artifacts. The signal processing chain of FE+NLC+ASAR provides the ability to sense neural activity concurrent with stimulation. Each of the steps in this chain can be configured and included/bypassed in the pipeline. The Sense IC provides a three-wire SPI interface. It also down-streams the commands to control the Stim IC. The controller integrated into the Sense IC implements the state machine for serial peripheral interface (SPI) communication, schedules the data for the sensing output, and features the capability of individual control of every FE/NLC/ASAR module [33–36]. The phase-locked stimulation (PLS) IC is a previously developed digital chip that supports 16-channel detection of the power at selectable frequencies within theta band (3–8 Hz), and triggers configured stimulation at a specified phase of the detected oscillation [41–42].

We designed a layout and manufactured a digital (Fig. 1b-middle) and an analog printed circuit board (PCB, Fig. 1b-bottom) using specialized software (Altium Designer 14.0) where each board consisted of 2 PCB layers. The Sense, Stim, and Spike IC footprints were placed on the analog layer and the PLS IC footprint on the digital layer. The SPI interface was routed from the analog layer input/output connector to the Sense IC and from the Sense IC to the Stim IC (Fig. 1b-right). We used a SPI with 3 wires: clock, master input/output slave (MISO), and master output/input slave (MOSI). Two-wire shift register interfaces were routed from the analog/digital layer input/output connector to the PLS IC/Spike IC (Fig. 1b-right). The sensing and stimulation FEs were routed to the two Omnetics PS1-16-AA connectors to which electrodes are connected. The digital and analog layer input/output connectors are compatible and can be stacked on top of each other. On the top connector, we placed the Xilinx Spartan 6 (XC6SLX150-2FGG484C) FPGA board to serve the role of the communication layer (Fig. 1a-top). The FPGA is configured to support four SPI interfaces and five shift registers, thus allowing up to four analog layers to be stacked together. We used a two-analog layer setup for all in-vitro and in-vivo experiments. Since we used separate SPI interfaces for each analog layer IC, the 4^th^ wire (select) on the SPI was not needed in the PCB design. The FPGA contains a finite state machine (FSM) that converts USB input (FTDI controller) into SPI (SPI controller) packet stream and vice-versa. For FPGA programming, we used the Xilinx ISE 14.2 software. Briefly, The FSM always begins with a Reset state after a reboot, and then enters an Idle state in which it waits for incoming packets. Once a packet is available, the FSM receives it byte by byte (Receive Byte) until the complete message is transferred (Receive Packet). The received packet is then being processed (Process Packet), converted into the appropriate interface (e.g., USB to SPI), and transmitted to the Neuro-stack ICs (via SPI or Shift Register). Similarly, after the processing is done, the response packet from the ICs enters a state during which it can transmit the packet (Transmit Packet) byte by byte (Transmit Byte) externally. Once the transmission is done, the FSM goes back to the Idle state and waits for new packets unless the streaming of the neural data is taking place, in which case the FSM enters Process Packet state indefinitely until the recording is stopped (Fig. S3-left). Stacked layers were placed inside a plastic enclosure (Fig. 1a) and wrapped from the inside with copper foil shielding tape to reduce the impact of the noise. Custom headstages (Fig. 1b, d) were built on a protoboard by placing two 5 × 2 connectors on each, which were internally routed to the Omnetics connector.

Neuro-stack’s communication layer uses a USB interface for external connections and a specific communication protocol that can address, configure, and start/stop each IC. The protocol is described by a packet structure (up to 520 bytes) that captures Command (such as Reset, Start/Stop, Read/Write configuration registers, etc.), Board ID (to select analog layer), Spike and PLS commands, and optional Payload (varies in length [Payload Length] depending on the command). The FPGA’s FSM processes the input packet and decides which IC is to be addressed and forwards relevant bytes to it. The protocol also includes safety error and cyclic redundancy check bytes (Fig S3-bottom). Every command returns its specific acknowledgment receipt indicating that the execution of the command was successful.

### Neuro-stack Software Design

The Neuro-stack graphical user interface (GUI, Fig. 1a) was built as a Universal Windows Platform application using Visual Studio (2017) and the Visual C# language. The application can be installed on any Windows (8.1 or higher) machine. We specifically used Surface Pro 5 for running the GUI application. The application uses a USB connection to directly communicate with the Neuro-stack (Fig. 1a) to enable viewing and configuration of real-time neural data, the configuration of PLS and other stimulation parameters, and manually triggered delivery of stimulation.

As an alternative to the GUI, the Neuro-stack application programming interface (API) is a library of functions built-in C++ that the user can call in custom-design experiments. The API combines all core and backhand GUI functions into a faster and more resource-efficient implementation. It’s built as a multi-thread real-time software pipeline, which threads mirror hardware blocks (e.g., Sense Process controls the Sense IC, Stim Process controls the Stim IC, etc. [Fig. S3-middle]). Processes responsible for each IC run in parallel and asynchronously forward commands to their associated IC or they await a command receipt or a recorded neural sample via the Input Queue (Fig. S3-middle). Neural samples are timestamped using network time protocol [NTP, 52] in the Sense and Spike Process threads upon their arrival. They are sent together with a sample value either to an external device or stored in Log Memory (Fig. S3-middle), which was used for synchronization. The library can be compiled for commonly used Linux, Windows, macOS, or ARM based target devices. We used the ARM-based (NXP i.MX 8M SoC) Coral Development Board to run the Neuro-stack API. To utilize all Coral Development Board capabilities, we complemented the library with functions that can store/save the TensorFlow Lite model and run inference on recorded neural samples using the Coral dev Board’s onboard tensor processing unit (TPU). Coral dev Board supports both wired (USB-C) and wireless (using a local network access point and a TCP/IP server with a X.509 certificate authentication) interfaces with external control capability and use of a real-time monitoring device (e.g., Experimental Computer). X.509 is a digital certificate that uses public key infrastructure. We used self-signed certificates since we only used one Experimental Computer to connect to the Neuro-stack. We used a MacBook Pro (2015) laptop as an Experimental Computer, which ran a client Python 3.6.9 script for triggering sensing, stimulation, TPU-specific commands, and transferring/storing/monitoring neural activity by using the Neuro-stack API running on the Coral dev Board (Fig. S3).

For in-vivo resting state neural recording experiments, we used the GUI application to control the Neuro-stack (Fig. 1). For in-vitro testing, in-vivo macro-stimulation (Fig. 3), behavioral stationary (Fig. 4) and ambulatory experiments (Fig. 2), we used the Neuro-stack API and Coral dev Board wireless configuration (Fig S3).

### Neuro-stack in-vitro testing

In-vitro studies involved the use of an oscilloscope, a phosphate-buffered saline (PBS) solution, a National Instruments digital to analog converter (NI-DAC), and the Neuro-stack (using both wired and wireless configurations; Fig. S4). Testing of the Sense and Spike ICs involved feeding 100 s of pre-recorded LFP/single-unit data through the NI-DAC. The analog signals were observed using an oscilloscope and recorded by a single channel using the Neuro-stack. For visualizing results, a time domain comparison was used for Sense IC and Spike IC (Fig. S5). The Stim IC was tested as part of closed-loop delay measurements and in previous reports [34]. Delivered stimulation was captured by the oscilloscope and on one channel using the Neuro-stack (Fig. S4, S5). The PLS IC was tested in-vitro as part of a previous study [41–42].

The round-trip delays were measured by sending a pulse train (50 pulses, 20 mV amplitude, 1 s pulse width, duty cycle 50%) from the NI-DAC to one channel recorded using the Neuro-stack. The modified software on the Coral dev Board continuously pooled incoming samples and detected the increase from zero (rising edge) in these incoming values. Once detected the rising edge triggered one-pulse of stimulation. The delay (mean ± standard deviation [std] for 50 pulses) was measured on the oscilloscope by capturing both the recording input and stimulation output rising edges and their time difference (Fig. S5d).

The Neuro-stack system and software latency from the recording input to the Sense Process thread on the Coral dev Board was measured using the same pulse train process but instead of triggering stimulation, the detected rising edge triggers a 1 s pulse to the Coral dev Board general-purpose input/output (GPIO) pin. We used the oscilloscope to observe the recording input and GPIO output, and measure the time difference between the rising edges (Fig. S5), which was equivalent to the system latency (mean ± std for 50 pulses).

### Neuro-stack in-vivo testing

#### Participants

Research participants were 12 patients (mean age 24.15 years, 9 females) with pharmacoresistant epilepsy who were previously implanted with acute stereo EEG depth electrodes for seizure monitoring. Participants volunteered for the research study during their hospital stay by providing informed consent according to a research protocol approved by the UCLA IRB. In each patient, 8-12 flexible polyurethane depth electrodes (1.25 mm diameter) were implanted solely for clinical purposes and prior to completion of the research study. Each depth electrode terminated in a set of eight insulated 40-μm platinum-iridium microwires (impedances 200-500 kΩ).

#### Electrode Localization

Electrodes were localized to specific brain regions using methods that have been previously used [53]. Briefly, a high-resolution post-operative CT scan was co-registered to a pre-operative whole brain MRI and high-resolution MRI using BrainLab stereotactic localization software (www.brainlab.com and FSL FLIRT (FMRIB’s Linear Registration Tool [54]). Medial temporal lobe (MTL) regions, including the hippocampus and entorhinal cortex, were delineated using the Automatic Segmentation of Hippocampal Subfields (ASHS [55]) software using boundaries determined from MRI visible landmarks that correlate with underlying cellular histology. White matter and cerebral spinal fluid areas were outlined using FSL FAST software [56]. Macro- and micro-electrode contacts were identified and outlined on the post-operative CT. For a list of localized brain regions in all participants see Table S1.

### Data Acquisition and Stimulation

For all in-vivo validation sessions, a Neuro-stack with two analog layers was used, which allowed for up to two micro-electrode bundles (16 channels) and eight macro-electrodes (16 bipolar channels). All micro- and macro-electrode recording sessions were sampled at 38.6 kHz and 6250 Hz, respectively. Base recordings were done without hardware decimation, non-linear correction, and artifact rejection on the Sense IC. Refer to *Data Analysis and Statistics* section for details about data analyses.

Macro-stimulation was performed in three participants while they rested in their hospital beds. In the first two participants, three stimulation bursts (0.5 mA) were delivered to a single bipolar electrode channel. In a third participant, we performed stimulation propagation mapping, where macro-stimulation was delivered to a single bipolar channel (Fig. 3c-d) and recording was done in the other 32 channels (Fig. 3e,h,i). During macro-stimulation, signal propagation was observed with using the following stimulation parameters: Channels: 1 out of 128; Amplitude: 0.25, 0.5, 0.75, 1.00, and 1.25 mA; Frequency: 60, 80, 100, 120, and 140 Hz; Pulse Width: 1.28 ms; Interphase Width: 150 us; Polarity: Anodic; Shape: Rectangular; Interburst delay: 16.67 s. The desired burst frequency was achieved by setting the inter-pulse delay appropriately.

Rectangular pulses recorded in all 32 channels were identified by using cross-correlation across all channels against a template waveform of the delivered stimulation pulse, which was later used for alignment (Fig. 3f,g) and calculating statistics of propagation with respect to varying amplitudes (Fig. 3h) and frequencies (Fig. 3i). For statistical calculations of the propagated power, all pulse waveforms across channels were normalized using the same value of the largest pulse that was propagated.

### Ambulatory Walking Task

Single-unit data was recorded in six participants during an ambulatory walking task. Two of the participants were instructed to walk around their hospital room freely and visit prominent ‘landmarks’ such as locations near windows, doors, tables, etc. A separate group of four participants was instructed to walk repeatedly (10 times) from one position to another position in the room using a linear path (Fig. S2b). The ambulatory movement was tracked using an eye-tracking headset (Pupil Labs Core device [57]) which contained inward-facing eye cameras (sampling rate: 200 frames per s) and an outward-facing world-view camera (sampling rate: 120 frames per s). Neuro-stack was connected to two micro-wire electrode bundles (Behnke-Fried, Ad-Tech) to record from 18 micro-wire contacts (16 recorded single-unit activity and 2 served as reference contacts). Recordings with respect to local references (same bundle) were recorded at a sampling rate of 38.6 kHz.

During the walking task, the participants wore an eye-tracker headset and a small backpack (Fig. 2a), which carried the Neuro-stack, the TPU (Coral dev Board) using the wireless configuration (Fig. S3), and a Voltaic V75 USB Battery Pack. The researcher used an Experimental Computer running an application (Python) to start/stop recordings and view in real-time the neural data. Both the Neuro-stack and eye-tracker were connected to the same local network from which the NTP timestamps were fetched. For a redundant method of synchronization, a miniature LED was attached to the corner of the world-view camera on the eye-tracking headset (Fig. 2a, Fig. S2d). The LED was programmed to turn on for 50 ms every 20 s during the experimental walking task, which was not visible by the participant and was also NTP-timestamped.

### Stationary Verbal Memory Task

Verbal memory performance was decoded using the Neuro-stack in a single participant. The memory task began with an encoding period, where the participant was instructed to learn a list of 10 words that were randomly selected and serially presented in an audio and visual format on an iPad Pro (3^rd^ generation) screen (Fig. S3 – top right). During encoding, each word was presented for 2 s with an inter-trial fixation period of 4 s. Words were drawn from clusters of six and seven of the word norms and were all 4-8 letter nouns that were rated as highly familiar (range 5.5-7 on a 1-7 scale), moderate to high on concreteness and imagery (range 4.5-6 on a 1-7 scale), and moderate in pleasantness (range 2.5-5.5 on a 1-7 scale) [58]. After the encoding period, participants completed a distractor task where they were instructed to determine whether a presented number (1-9) was Odd or Even. The distractor task was then immediately followed by a verbal recall period where participants were cued to verbalize as many words as they could remember during a 30 s period. During the experimental paradigm, encoding, distractor, and retrieval periods were repeated 10 times. Memory performance was calculated as the proportion of previously encoded verbalized words that were recalled.

During the verbal memory task, we used the Neuro-stack in a wireless configuration (Fig S3) together with both the Experimental Computer and Stimulus Presentation device (iPad). We used the Sense IC to record 16 channels from two (left/right hippocampus) micro-wire bundles. Stimulus presentation on the iPad was implemented as a game using Xcode 11.2.1 and Swift 5.0.1 programming languages. For network communication, we used two TCP (transmission control protocol) channels (Fig. 4b, Fig. S3; 1. Experimental Computer – Coral dev Board, 2. Experimental Computer – iPad). For online binary classification of the incoming neural data into remembered/forgotten words, we used a pretrained neural network model (2 × CNN1D + LSTM + Dense; Fig. 4e). The background processing of the task’s data was divided into two phases: 1) training and 2) prediction, consisted of 5 and 4 blocks of the verbal memory task cycle, respectively (Fig. 4c, presented 7 blocks only; 3 training and 4 prediction). The purpose of the training phase was to personalize the model for the participant. Only the last two Dense layers from the model were used for retraining and embedding selected filters into the prediction model. The training phase involved downsampling and filtering of raw data (0.1 – 250 Hz), packing the data separately for each observed brain region (Preprocess step), and transmitting packages from the Neuro-stack externally to the Experimental Computer where the model retraining took place (Fig. 4b). The words were presented using an iPad Pro tablet, which also used a built-in speech recognition algorithm to supply real-time outcomes (i.e., remembered or forgotten) to the Experimental Computer. The word onset events were isolated and weighted using a Gaussian window where one standard deviation was 2.5 s and cutoffs were made at −5 and 5 s (before and after word onset), thus giving data around the word onset higher priority. The retraining of the model took place during every Distraction phase (30 s) of the verbal memory task. Once retrained, the model was automatically converted on the Experimental Computer from Tensorflow 2.2 to Tensorflow Lite and uploaded wirelessly to the Edge TPU. During the prediction phase, the same format of preprocessed data was rerouted to the Edge TPU, where prediction took place. The predictions from TPU and labels from the iPad were transmitted to the Experimental Computer for performance assessment after each word trial (Fig. 4b).

#### Neural Network Model

The neural network model (Fig. 4e) was used to decode performance on the verbal memory task in a single participant. The model architecture included two one-dimensional convolutional neural networks (CNN1D) (1st with 32 nodes and 2^nd^ with 64 nodes) and a long-short term memory (LSTM) neural network layer with 64 nodes. The L2 regularization was used in the CNN1D and Dense layers and was proportional to the square of the weight coefficients’ value. Moreover, the training dropout technique [59] was applied after each layer with a 0.2 rate, except for the LSTM, which used a 0.1 rate and a recurrent dropout (0.5 rate). The complete structure of one branch is presented in Fig. 4e. The branches were structurally identical for all brain regions but had different weights after training. The model was pretrained offline using data from 6 medial temporal lobe regions (left/right anterior hippocampus, left/right posterior hippocampus, left/right entorhinal cortex) from 10 participants who performed the exact same verbal memory task (Fig. 4a) previously using a Blackrock Neuroport system to record neural data. LFP data (sampling rate 250 Hz, batch size 512) was extracted around the verbal memory task word onsets (same Gaussian window as before) and fed into the model for training (Fig. S6a). The data from all participants was divided into training (50%), validation (25%), and test (25%) sets. Then training and validation datasets were combined, shuffled, and used for training of the base model (Fig. S6c). Binary cross-entropy was used for the loss function, with root mean square propagation for the optimizer (learning rate of 0.001). Five-fold cross-validation (Fig. S6d – average across folds) was used for validation using the presented hyperparameters. Hyperparameter optimization of the final decoding model (Fig. 4e) was done during the validation phase and with respect to the F-1 score (0.5 threshold). During the training phase with Neuro-stack, we used the same training parameters except that CNN1D and LSTM layer coefficients were fixed and only Dense coefficients were adjusted. Also, we only used two model branches out of six that were previously trained on the Blackrock-acquired data (hippocampal channels only) to match the left/right hippocampal electrode placement in the single participant who performed the verbal memory task Neuro-stack experiment. During online training phase, all incoming windows of the LFP data were continuously combined with the previous windows and used for retraining, while new retraining iteration updated coefficients saved from the previous retraining block. Participants (Blackrock: B1-B10; Neuro-stack: N1), their memory performance during verbal memory task, and test accuracies using offline (B1-B10) and online (N1) models are shown in Fig. S6b.

To isolate frequency bands that were the most significant for the neural network model decisions, we adapted Grad-CAM [45] for one-dimensional CNN and applied it on each branch separately. By doing this, we isolated activation filters of the second CNN1D (Fig. 4f,g – time-frequency representation).

The above described neural network model was chosen after an extensive trial and error process during which multiple classification algorithms were tested on the same dataset. Specifically, before utilizing the neural network model, the data was classified using shallow methods such as Support Vector Machine (SVM). As part of the feature engineering process, we supplied SVM models with raw, power, and phase data in 0-250 Hz range chunks of 7 s (word onset at 3.5 s) or in a sequence of 1 s sliding time windows (with no overlap). Before choosing the final decoding model, we also tested several convolutional and recurrent neural network (RNN) architectures. Summary of accuracies for each of these decoding methods is presented in Table S3).

### Data Analysis and Statistics

#### iEEG Power Spectrum Extraction

All time-frequency power scalograms were obtained using CWT (Continuous Wavelet Transform - MATLAB *cwt* command) performed on z-scored time domain data (each channel normalized separately). The base wavelet chosen was the complex Morlet with a symmetry parameter (gamma) equal to 3 and a time-bandwidth product equal to 60. The wavelet coefficients were calculated at seventy logarithmic frequency points from 1 to 125 Hz, after which the squared absolute value of the coefficients resulted in a power scalogram.

All frequency power spectrums were obtained using FFT (Fast-Fourier Transform - MATLAB *fft* command). The FFT length chosen was the largest power of 2, less than the length of the observed iEEG trace. The coefficients were then normalized with the trace length. Finally, the squared absolute value of the spectral coefficients multiplied by 2 (one-sided FFT) resulted in the power spectrum.

#### Spike sorting

We performed spike sorting using Wave_clus 3 [44]. Preprocessing included the use of a notch-filter to remove 60 Hz noise. Selected clusters were chosen so that more than 250 spikes were identified and that out of these, 1% or less had inter-spike-intervals (ISI) of less than 3 ms.

### Data and Code Availability

Data and code are available upon reasonable request.

