## Supplemental Information for "A wearable platform for closed-loop stimulation and recording of single-neuron and local field potential activity in freely-moving humans"

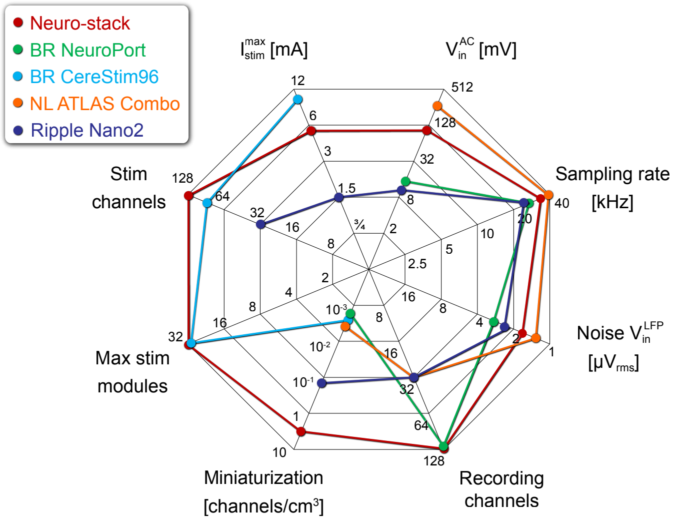


**Figure S1. Comparison of Neuro-stack with commonly used bedside intracranial recording and stimulation systems used in humans (Related to Figure 1)**

Neuro-stack capabilities as compared to existing human intracranial recording and stimulation systems. Characteristics shown include the device sampling rate, noise of the Sense IC (Noise $V_{in}^{LFP}$ ), number of recording channels that can be used, linear input dynamic range ($V_{in}^{AC}$ ), maximum stimulation current ($I_{stim}^{max}$), number of channels that can be used for stimulation (Stim channels), and total number of stimulation channels that can be used simultaneously (Max stim modules). The main advantages of the Neuro-stack come from the miniaturization of the electronics per channel (channels/cm$^3$) that allow for its small size and wearability and its integrated full-duplex capability that incorporates both stimulation and sensing (red line). BR: Blackrock, NL: Neuralynx.

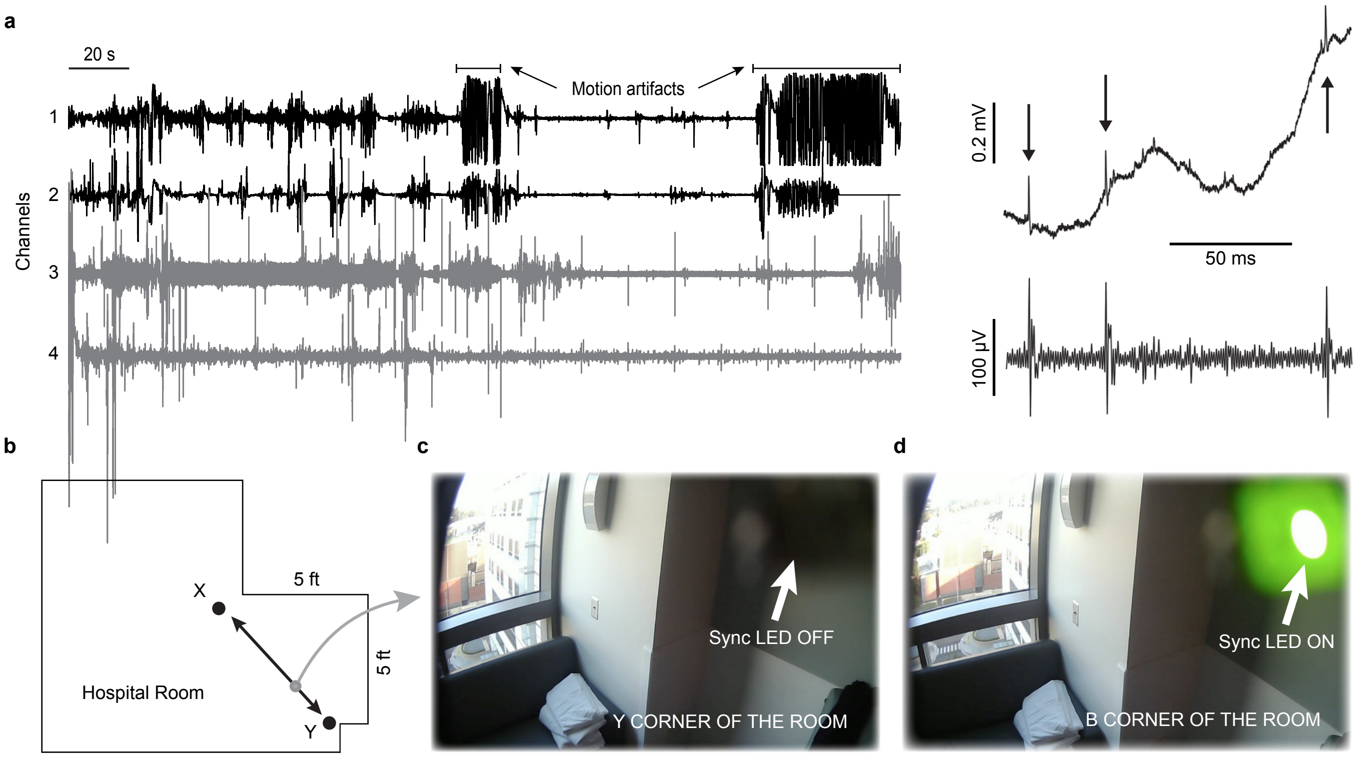


**Figure S2. Overview of the ambulatory walking task (Related to Figure 2)**

**a,** Motion artifacts from four example Neuro-stack recording channels from an example participant during walking behavior when referencing is done using a separate (black, channels 1-2) or the same (grey, channels 3-4) micro-wire bundle. Single-unit activity (example spikes in raw and filtered data shown to the right) that can be extracted using waveform shape and temporal differences as compared to the motion artifact.

**b,** Shown is a top-down view of the hospital room layout in which an example participant completed the walking task, during which they were asked to walk back and forth between points X and Y repeatedly. Points X and Y were placed within a small area of the hospital room (~25 ft$^2$).

**c,** Example screenshots from the world-view camera as an example participant approached point Y from X (gray arrow shown in **b** that was captured via an eye-tracking headset.

**d,** Same as **c** but a screenshot when the LED pulse that is used for synchronization (sync LED) was turned on versus off (**c**).

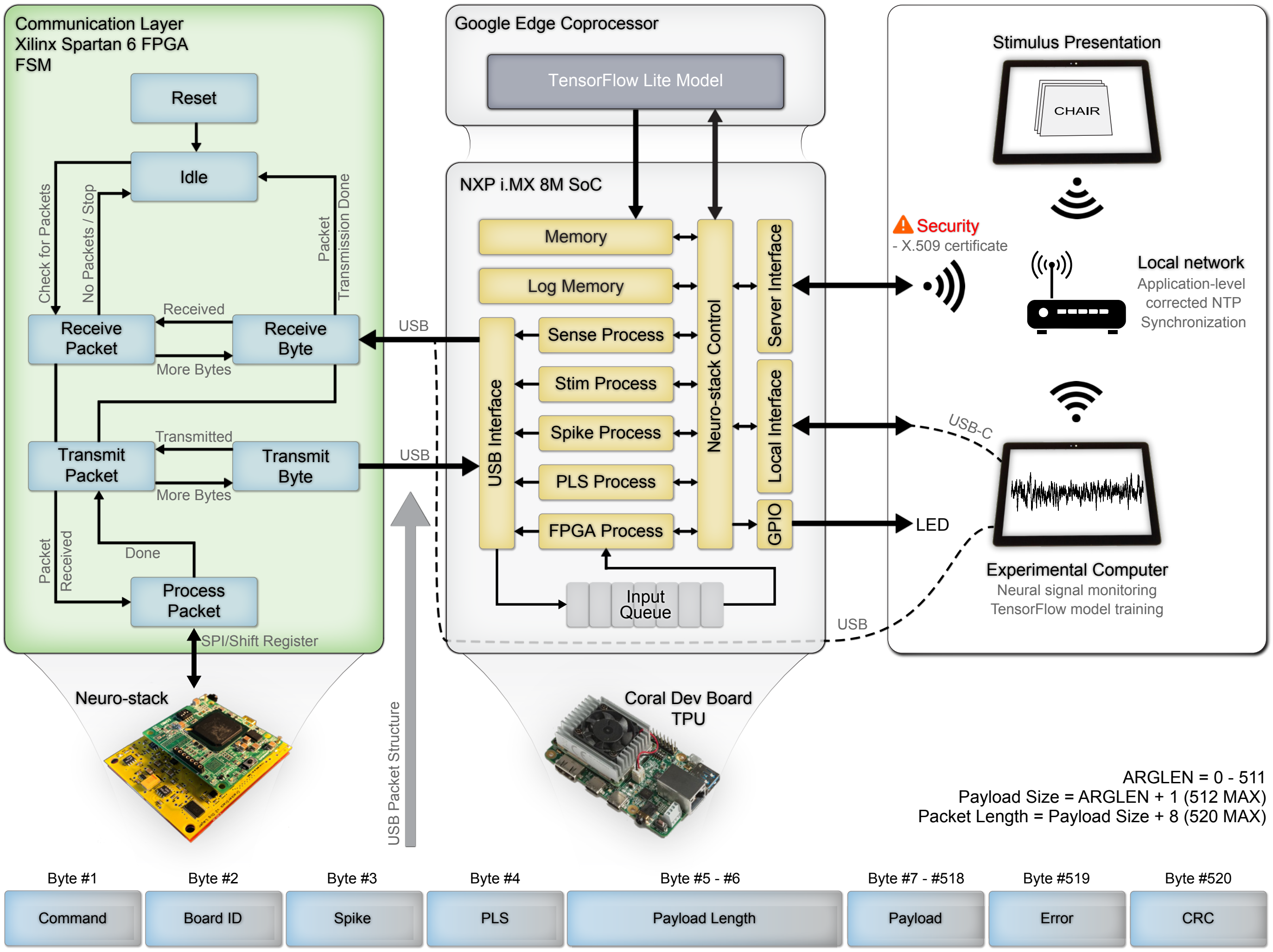

Communication Layer
Xilinx Spartan 6 FPGA
FSM
Reset
Idle
Check for Packets
No Packets / Stop
Packet Transmission Done
Received
Receive Packet
Receive Byte
More Bytes
Transmitted
Transmit Packet
Transmit Byte
More Bytes
Packet Received
Done
Process Packet
SPI/Shift Register
Neuro-stack
USB
USB
USB Packet Structure
Google Edge Coprocessor
TensorFlow Lite Model
NXP i.MX 8M SoC
Memory
Log Memory
Sense Process
Stim Process
Spike Process
PLS Process
FPGA Process
USB Interface
Neuro-stack Control
Server Interface
Local Interface
GPIO
Input Queue
Coral Dev Board
TPU
USB
Stimulus Presentation
CHAIR
Security
- X.509 certificate
Local network
Application-level corrected NTP Synchronization
USB-C
LED
Experimental Computer
Neural signal monitoring
TensorFlow model training
ARGLEN = 0 - 511
Payload Size = ARGLEN + 1 (512 MAX)
Packet Length = Payload Size + 8 (520 MAX)
Byte #1
Command
Byte #2
Board ID
Byte #3
Spike
Byte #4
PLS
Byte #5 - #6
Payload Length
Byte #7 - #518
Payload
Byte #519
Error
Byte #520
CRC

**Figure S3. Neuro-stack experimental setup (Related to Figure 1-4)**

A simplified block diagram showing Neuro-stack setup including the (LEFT) Communication Layer, which uses a field-programmable gate array (FPGA, Xilinx Spartan 6 model) with a finite-state machine (FSM) built within it to allow for receipt of external messages (packets) via USB (e.g., stimulation command) and streaming of the neural data. These packets are then processed and converted to serial peripheral interface (SPI) or Shift Register packets, and transmitted to the Neuro-stack integrated circuits (ICs, SPI: Stim and Sense IC; Shift Register: Spike and PLS IC). This FSM is bi-directional and thus also processes SPI packets received from the Neuro-stack and converts them into USB packets, which are then transmitted to the external Coral Development Board (Coral Dev Board) device. The FSM always begins with a Reset state after a reboot, and then enters an Idle state in which it waits for incoming packets. Once a packet is available, the FSM receives it byte by byte (Receive Byte) until the complete message is transferred (Receive Packet). The received packet is then being processed (Process Packet), converted into the appropriate interface (e.g., USB to SPI), and transmitted to the Neuro-stack ICs (via SPI or Shift Register). Similarly, after the processing is done, the response packet from the ICs enters a state during which it can transmit the packet (Transmit Packet) byte by byte (Transmit Byte) externally. Once the transmission is done, the FSM goes back to the Idle state and waits for new packets unless the streaming of the neural data is taking place, in which case the FSM enters Process Packet state indefinitely until the recording is stopped. (CENTER) Coral Dev Board that can directly communicate with the Neuro-stack Communication Layer (via USB connection) using an Application Programming Interface (API, shown here) or a device (e.g., Experimental Computer) with an installed GUI via USB (Fig. 1a). The Coral Dev Board contains a regular ARM-based central processing unit (NXP i.MX 8M SoC) and a Google Edge Coprocessor, a tensor processing unit (TPU). The Neuro-stack API library runs on the ARM processor and contains a real-time pipeline for handling control and data flow to and from the Neuro-stack for each IC and Communication Layer (or FPGA). The Input Queue handles streams of both neural data and acknowledgment receipts from the Communication Layer and redirects them to the appropriate block on the Coral Dev Board responsible for each ICs (e.g., Sense, Stim, … Process). The Neuro-stack Control block contains all of the API functions, which are then multiplexed to additional layers responsible for wireless (via Server Interface) or wired (via Local Interface) communication with the Experimental Computer. Additionally, the Neuro-stack Control block also contained functions for controlling the TPU, such as loading/saving the machine learning model (TensorFlow Lite Model) to/from the Memory block, redirecting the data streams directly towards the TPU, and receiving the TPU's output once it is ready. The incoming neural data streams can also be stored locally in Log Memory or transferred to external storage on the Experimental Computer through the Neuro-stack Control block. Furthermore, an LED light can be triggered (to turn on/off) through available general-purpose input/output (GPIO) pins for synchronization purposes. These triggered on/off events are internally temporally aligned with the incoming neural data in order to synchronize it with data from eye-tracking cameras. (RIGHT) A local network can be created either by using a separate access point (shown here, e.g., router, hotspot, etc.), or by the Coral Dev Board, which contains a network controller that can support access point topology and thus can create its own local network. This wireless mode means that a server is created on the Coral Dev Board to allow for other devices, such as the Stimulus Presentation device (e.g., iPad used to present the verbal memory task) or Experimental Computer (e.g., to view neural data in real-time) to access the Neuro-stack API functions. Security warning points to the importance of a safe wireless connection, implemented by X.509 certificate authentication. Wired mode is also supported through Local Interface block (e.g., Experimental Computer connected via USB-C). All devices connected to the local network use Network Time Protocol (NTP) to log events with timestamps fetched from a common server in order to synchronize them. (BOTTOM) The structure of the USB packets sent from the Coral Dev Board to the Neuro-stack Communication Layer, which contains up to 520 bytes that describe the type of Command, Board ID to address specific analog layer, Spike byte, phase-locked stimulation (PLS) byte, and Payload for additional information where its length (Payload Length) depends on the type of command. The packet also contains bytes for error codes (Error) and a cyclic redundancy check (CRC) to detect accidental changes in the raw packets. The communication layer extracts a relevant portion of the USB packet, converts it to desired interface (SPI or Shift Register), and transmits it to the addressed IC.

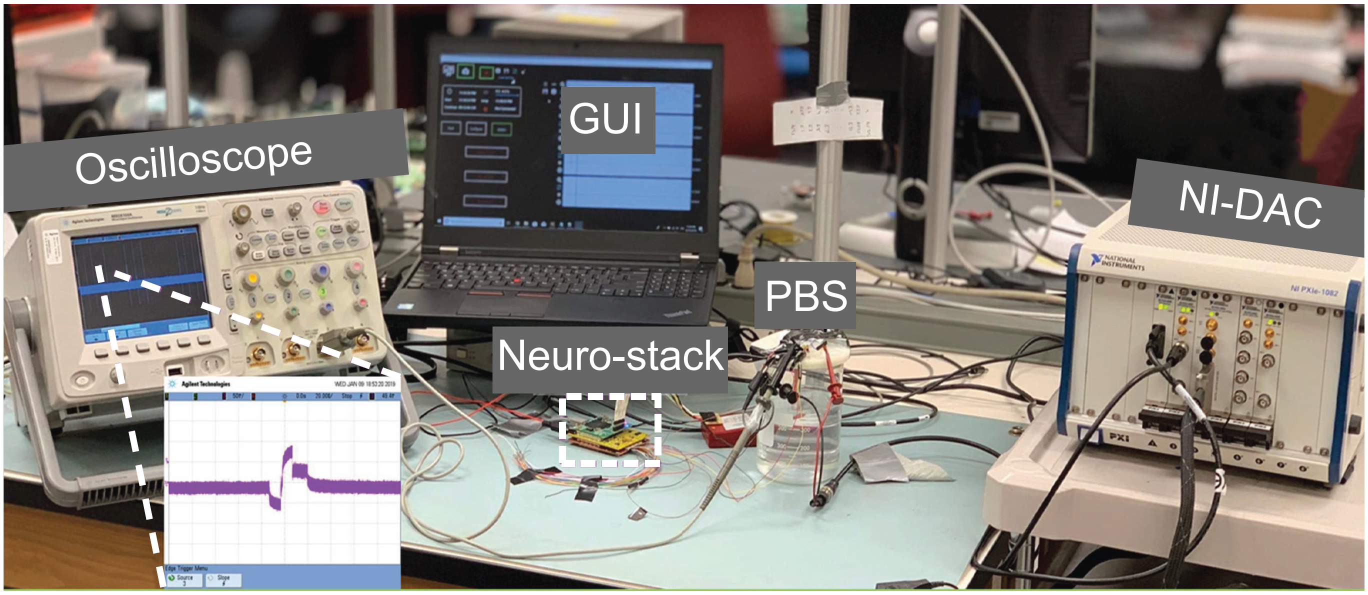


**Figure S4. Neuro-stack in-vitro validation setup (Related to Figure 1)**

Setup for in-vitro validation of recordings and stimulation includes the Neuro-stack platform and GUI, a phosphate-buffered saline (PBS) solution, a NI-DAC (National Instruments Digital to Analog Converter) device for conversion of pre-recorded neural signals, and an oscilloscope for monitoring the signal and measuring delivery time (of synchronization and stimulation pulses). The oscilloscope shows a single pulse of stimulation that was delivered using the Neuro-stack. Adapted from Alzuhair, 2019 [30].

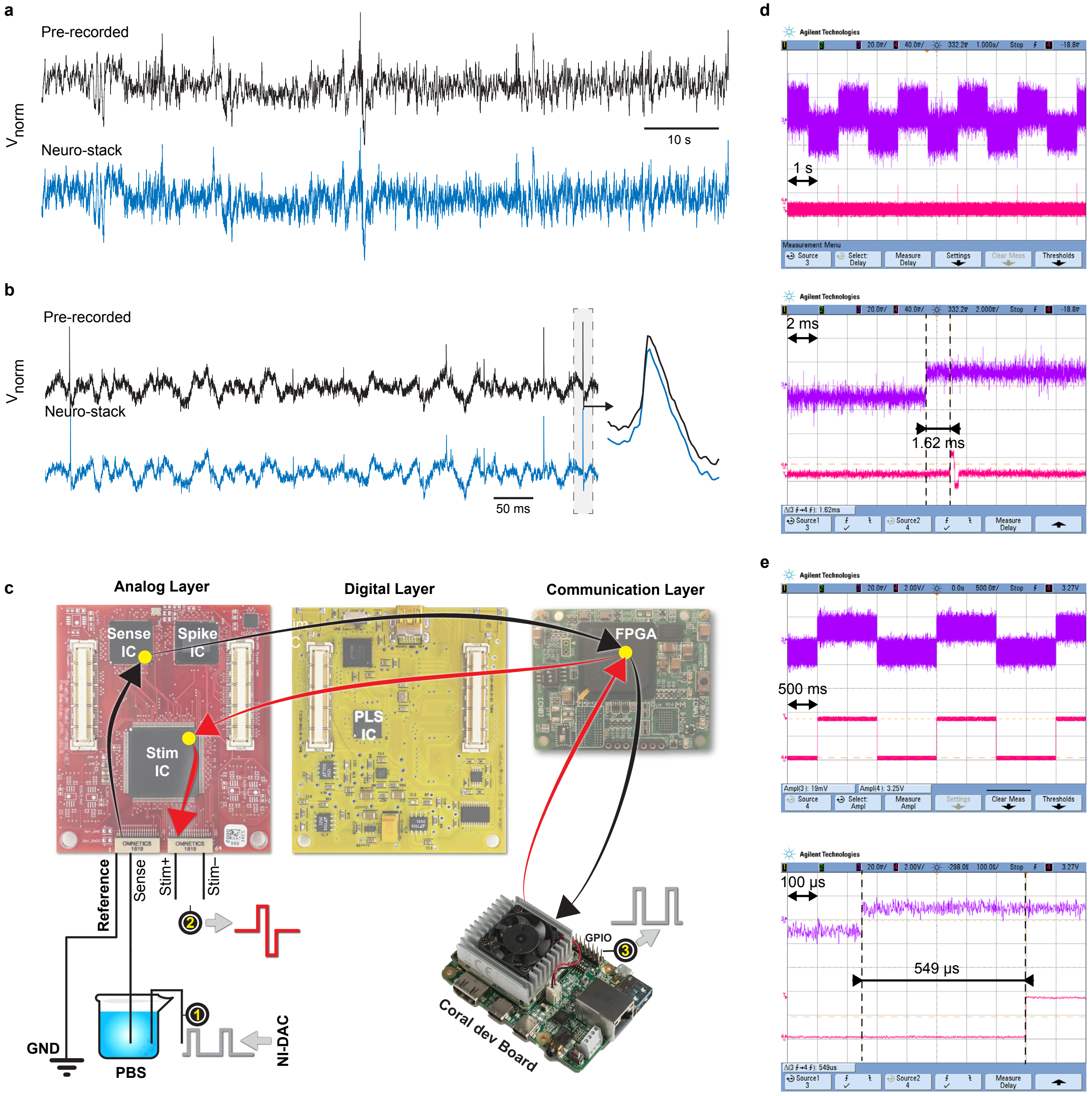


**Figure S5. Neuro-stack in-vitro validation results**

**a,** Pre-recorded and normalized LFP signal (top) fed into the Sense IC front-end and the resulting Sense IC recording (bottom). **b,** Pre-recorded and normalized single-unit signal (top) fed into the Spike IC front-end and resulting Spike IC recording (bottom). On the right is a zoomed-in comparison of a single spike waveform. **c,** Validation setup from Fig. S4 in more detail, showing signal path for in-vitro sensing (black arrows; NI-DAC - PBS - Sense IC - FPGA - CDB) and for in-vitro stimulation (red arrows; CDB - FPGA - Stim IC - PBS). The round-trip delay was measured from point 1 (signal generator) to the CDB (black arrows), and back (red arrows) to point 2 (stimulation output). Test sensing signal was a pulse train fed into the PBS, which triggers stimulation once detected on the CDB. The sensing system delay was measured from point 1 (signal generator) to point 2 (GPIO output). Test signal was again a pulse train, which triggered the GPIO pulse once detected on the CDB. The output stimulation current was converted to voltage using resistors and was observed on the oscilloscope. **d,** The round-trip delay observed on the oscilloscope (Point 1 and 2; Zoomed-in at the bottom). **e,** The system sensing delay observed on the oscilloscope (Point 1 and 3; Zoomed-in at the bottom).

a

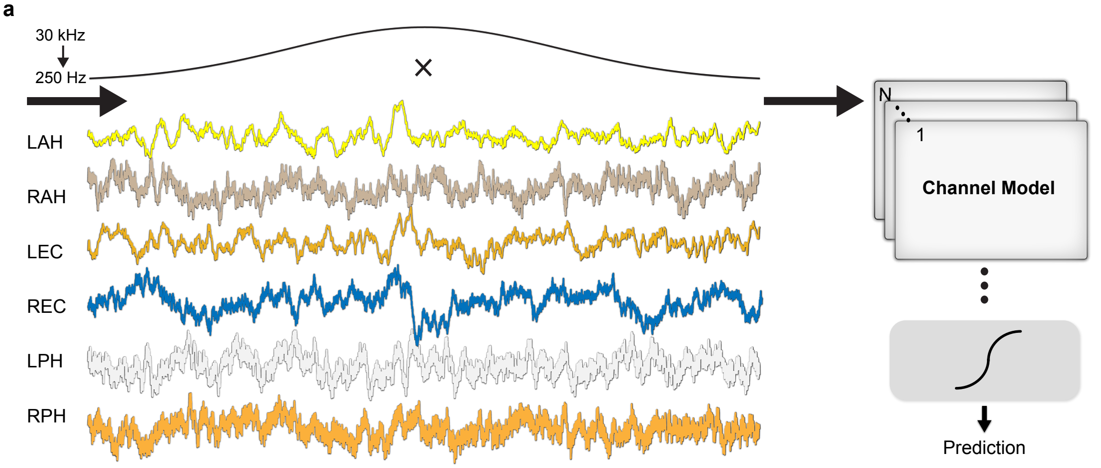


b

| Participant | Memory Performance [%] | Test Accuracy [%] |
|---|---|---|
| 1 | 40.0 | 92.96 |
| 2 | 44.8 | 88.49 |
| 3 | 56.4 | 91.71 |
| 4 | 49.1 | 88.35 |
| 5 | 28.3 | 92.08 |
| 6 | 17.7 | 90.22 |
| 7 | 30.0 | 99.50 |
| 8 | 32.0 | 91.45 |
| 9 | 31.6 | 88.80 |
| 10 | 28.7 | 93.50 |
| N1 | 38.9 | 72.5 |

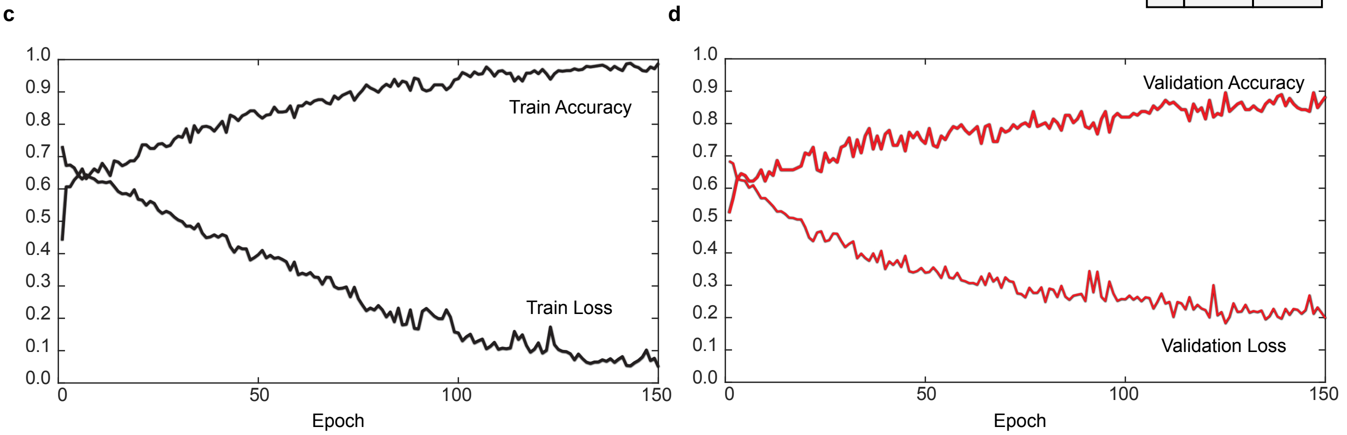


**Figure S6. Offline neural network model (Related to Figure 4)**

**a,** Downsampled (30 kHz to 250 Hz) windows of LFP data from 6 brain regions: Left/Right Anterior Hippocampus (LAH/RAH), Left/Right Entorhinal Cortex (LEC/REC), and Left/Right Posterior Hippocampus (LPH/RPH) were range [-1,1] normalized and multiplied with a Gaussian, centered around the word onset. Each channel was then fed into a dedicated Channel Model (N = 6) and concatenated results were then classified. Note that data from all channels on one micro-wire were merged together and then divided into 6 brain region categories. Data was then split into training, validation, and test data sets for each participants (50, 25, 25 %). Training and validation data sets from all participants were merged, shuffled and used for training and 5-fold validation of the base model. **b,** 10 participants recorded using a Blackrock system and a single participant recorded using the Neuro-stack (N1). Shown are total memory performances for each participant including both training and test samples, and accuracy of the base model on test dataset for each participant (1-10). Test accuracy for participant N1 represented the result from using the online model, retrained in real-time during the experiment. **c,** Training accuracy and loss of the base model. **d,** Validation accuracy and loss, averaged across 5 folds.

| Participants | Brain Region | Macro-recording | Micro-recording | Macro-stimulation | Walking Task | Verbal Memory Task |
|---|---|---|---|---|---|---|
| 1 | Left Hippocampus | ✓ | | | | |
| 2 | Left Hippocampus | ✓ | ✓ | | | |
| 3 | Left Temporo-Parieto-Occipital | ✓ | ✓ | | | |
| 4 | Right Orbitofrontal | ✓ | ✓ | ✓ | | |
| 5 | Left Entorhinal<br>Left Hippocampus | ✓ | ✓ | ✓ | | |
| 6 | Left Hippocampus<br>Right Hippocampus | ✓ | ✓ | | ✓ | |
| 7 | Left Hippocampus<br>Right Hippocampus | ✓ | ✓ | | ✓ | ✓ |
| 8 | Left Hippocampus<br>Right Hippocampus | ✓ | ✓ | | ✓ | |
| 9 | Left Hippocampus<br>Right Hippocampus | ✓ | ✓ | | ✓ | |
| 10 | Left Hippocampus | ✓ | | ✓ | | |
| 11 | Left Hippocampus<br>Anterior Cingulate | ✓ | ✓ | | ✓ | |
| 12 | Left Hippocampus<br>Right Entorhinal | ✓ | ✓ | | ✓ | |

**Table S1. Participant’s electrode localization and types of conducted experiments (Related to Figure 1 - 4)**

For each participant, electrode localizations (brain regions) are shown as well as whether macro-recording, micro-recording, and/or macro-stimulation was done in conjunction with the walking and/or verbal memory task. Brain regions where electrodes were placed were based on clinical criteria and included the left/right hippocampus, left/right entorhinal cortex, anterior cingulate, left temporo-parieto-occipital junction, and right orbitofrontal cortex. A total of eight participants completed the walking task with micro- and macro-electrode recordings to capture single-unit and local field potential activity. The number of channels that were recorded ranged from 2 to 40 (mean: 13.85).

| | Blackrock Microsystems | Neuro-stack (4 analog layers) |
|---|---|---|
| **Model Name** | CereStim R96 Micro Stimulator | Neuro-stack Stim Engine |
| **Stimulation Channels** | 96 | 256 |
| **Stimulation Engines** | 3 | 32 |
| **Type of Protection** | Class II | Class II |
| **Degree of Protection** | Type BF Applied Part | Type BF Applied Part |
| **Output Voltage** | ±4.7 – ±9.5 V | ±6 V |
| **Polarity** | Selectable Anodic or Cathodic First | Selectable Anodic or Cathodic First |
| **Amplitude** | 1 μA – 215 μA, (1 μA) | 20 μA – 5,080 μA, (20 μA) |
| **Pulse Shape** | Rectangular | Custom (Rectangular/Triangle/Sine/Exponential) in 16 steps (8 bits) |
| **Frequency** | 4 Hz – 5 kHz | 2.37 Hz – 16.67 kHz |
| **Pulse Width** | 44 μs – 65,535 μs | 10 μs – 1,280 μs, (10/20/40/80 μs) |
| **Interphase Width** | 53 μs – 65,535 μs | Interphase: 0 μs – 150 μs, (10 μs)<br>Interpulse: 10 μs – 81.6 ms, (320 μs)<br>Interburst: 81.92 ms - 408.32 ms, (1.28 ms) |
| **Recommended Electrode Impedance** | < 100 kΩ | < 1 kΩ. High Impedance acceptable |
| **PC Hardware Interface** | USB A-B cable | USB mini B to type A cable or Wi-Fi |
| **Stim Manager PC Software Compatibility** | Windows 7 (32/64-bit) compatible | Windows 8.1 or higher |
| **API** | x86 & x64 library versions | Source and precompiled (ARM, Intel, AMD) |
| **Analog Resolution** | 12 bits | 12/21 bits |
| **External Power Supply** | PMP15M-13 Protek Power Supply<br>AC Input: 100 – 240 Vac, 0.5 – 0.3 A,<br>50 – 60 Hz DC Output: 15 V, 1 A MAX | DC Output Supplies ±6 V<br>USB 5 V |
| **Cable Connectors** | Samtec MIT-019-02-F-D | Omnetics PS1-16-AA and four 5 × 2 pin connectors per analog layer |
| **Monitor Connector** | Samtec SMM-109-02-F-D | Mini USB (data/power) or Wi-Fi |
| **Operating Environment** | 10 ˚C to 40 ˚C, 10 to 85% R.H. (non-condensing) | 10 ˚C to 40 ˚C |
| **Storage/Transportation Environment** | -15 ˚C to 60 ˚C, 10 to 85% R.H. (non-condensing), 500 to 1,060 hPA | -15 ˚C to 60 ˚C |

**Table S2. Comparison of the Neuro-stack and existing Blackrock CereStim stimulation engine (Related to Figure 1, 3)**
Shown are detailed features of stimulation engines. Major advantages of the Neuro-stack are the ability to customize stimulation waveforms (Pulse Shape, Interphase Width, etc.), a more flexible API (Application Programming Interface) library, and the ability for wireless control (PC Hardware Interface) for mobile experiments.

| # | Method | Features | Accuracy [%] |
|---|---|---|---|
| 1 | SVM | Power (TF) | 60 |
| 2 | SVM + TW | Power (TF) | 60 – 90 |
| 3 | SVM + TW + PCA | Power (TF)<br>Phase (TF) | 65 – 90 |
| 4 | GoogleNet + DENSE | Power (TF)<br>Phase (TF) | 69.40 – 96.87 |
| 5 | DENSE | Raw Data | 72.1 – 81.2 |
| 6 | CNN2D + DENSE | Power (TF)<br>Phase (TF) | 63.4 – 75.0 |
| 7 | LSTM/GRU + DENSE | Raw Data | 75.6 – 88.3 |
| 8 | CNN1D + DENSE | Raw Data | 72.1 – 87.9 |
| 9 | GP + DENSE | Raw Data | 40 – 60 |
| 10 | CNN1D + LSTM + DENSE | Raw Data | 88.3 – 99.5 |

**Table S3. Machine learning methods used for decoding memory using the Neuro-stack (Related to Figure 4)**

Accuracy comparison of the different algorithms used for decoding verbal memory performance from neural activity in real-time using the Neuro-stack. SVM: support vector machine; TW: time-windowing; PCA: principal component analysis; GoogleNet: a large Google's neural network, pretrained on time-frequency images of the electrocardiogram signals; Dense: fully connected neural network layer; CNN2D: 2D convolutional neural network layer; LSTM: long-short term memory neural network layer; GRU: gated recurrent unit neural network layer; CNN1D: 1D convolutional neural network layer; GP: gaussian process. Power (TF) and Phase (TF) refer to chunks of either power or phase time-frequency (TF) representations of the signal. Raw data refers to neural data that is minimally processed (downsampled) in the time-domain and fed directly into the decoding algorithm. See Online Methods section for more details.
